# DNA damaged-induced phosphorylation of a viral replicative DNA helicase results in inhibition of DNA replication through attenuation of helicase function

**DOI:** 10.1101/2023.09.29.560221

**Authors:** Caleb Homiski, Rama Dey-Rao, Shichen Shen, Jun Qu, Thomas Melendy

## Abstract

A major function of the DNA damage responses (DDRs) that act during the replicative phase of the cell cycle is to inhibit initiation and elongation of DNA replication. The polyomavirus SV40 is an important model system for studying human DNA replication and DDRs due to its heavy reliance on host factors for viral DNA replication, and the arrest of SV40 DNA replication in response to DDR activation. The inhibition of SV40 DNA replication following DDR activation is associated with enhanced DDR kinase phosphorylation of SV40 Large T-antigen (LT), the viral origin-binding protein and DNA helicase. NetPhos prediction of LT phosphorylation on multiple sites were confirmed by mass spectroscopy, including a highly conserved DDR kinase site, T518. In cell-based DNA replication assays expression of the phosphomimetic mutant form of LT at T518 (T518D) resulted in dramatically decreased levels of SV40 DNA replication; while LT-dependent transcriptional activation was unaffected. WT and LT T518D were subsequently expressed, purified, and analyzed *in vitro* for assessment of biochemical function. In concordance with the cell-based data, reactions using SV40 LT-T518D, but not T518A, showed dramatic inhibition of SV40 DNA replication. Importantly, the LT T518D mutation did not affect critical LT protein interactions or its ATPase function, but showed decreased helicase activity on long, but not very short, DNA templates. These results suggest that DDR phosphorylation at T518 inhibits SV40 DNA replication by impeding LT helicase activity, thereby slowing the DNA replication fork. This is consistent with the slowing of cellular replication forks following DDR and may provide a paradigm for another mechanism for how DNA replication forks can be slowed in response to DDR, by phosphorylation of DNA helicases.

## INTRODUCTION

Cellular DNA damage response (DDR) pathways play important roles in genome maintenance, including upregulating DNA repair pathways and inhibiting DNA replication so as not to exacerbate damage to the DNA during genome duplication (1-4). While many studies on DDR inhibition of DNA replication have focused on the inhibition of replication origin firing, it has also been shown that there is a nearly 10-fold decrease in replication fork elongation rate following activation of DDR in systems as diverse as yeast to human cells (5-7), as well as fork remodeling that allows for reversal of replication forks (8). Slowing of replication fork progression has been shown to be dependent on the apical DDR kinases, ATM, ATR, and DNA-PK, and works on forks distal to the sites of damage (Fig1A) (9-12). Recently one mechanism of replication fork slowing has been shown to be due to increased replication fork reversal (9); however, there have been some indications that modification of DNA replication fork proteins may also participate in replication fork slowing (13-16).

**Fig 1.**
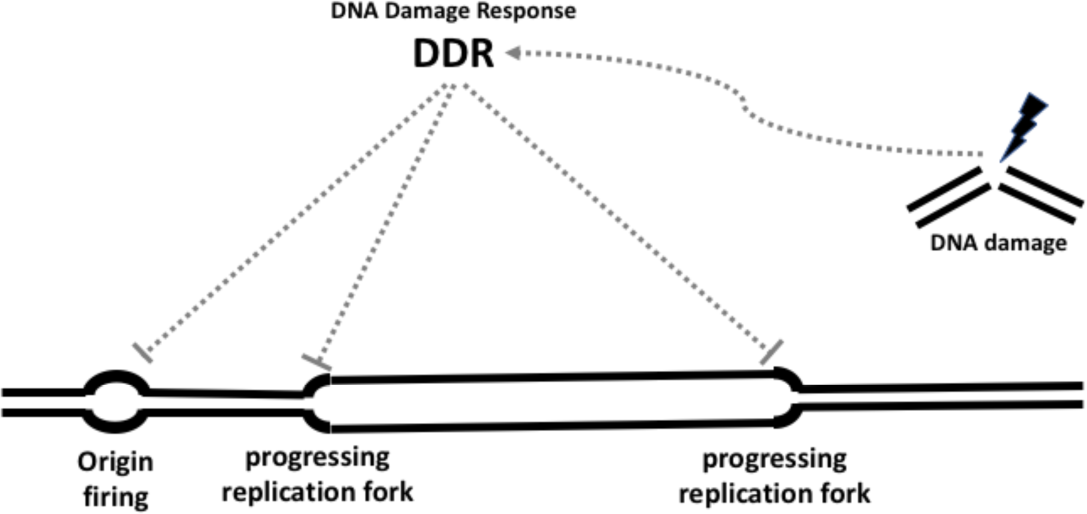
DNA damage inhibits both firing of origins and progression of existing replication forks in trans. DNA damage at distal sites, or even on separate DNA molecules (dark lines), can activate cellular DDR pathways (dashed gray lines) which can then inhibit the firing of late or subsequent DNA origins. DDR activation can also act to slow the progression of existing DNA replication forks by as much as 90%, even at replication forks that do not encounter DNA lesions. Both processes serve to slow DNA replication presumably to allow cellular DNA repair pathways opportunity to resolve the DNA damage prior to forks arriving at DNA lesions.

SV40 is a member of the polyomaviruses (PyVs), a family of non-enveloped viruses that have small circular double-stranded DNA (dsDNA) genomes, a single origin of replication (ori), and rely predominantly on the host cell DNA replication machinery to duplicate their genomes (17-19). SV40 has been used as a model system to investigate many human cellular processes, including the function and mechanism of several DNA replication factors (17-19). SV40 Large Tumor Antigen (LT) is the only viral protein required for replication of the SV40 genome. LT binds to the SV40 origin of replication (ori) and forms a double hexameric DNA helicase to initiate bidirectional replication from the ori (17-19). The LT hexamers directly recruit host DNA replication factors including Replication Protein A (RPA), Topoisomerase I (TopoI), and Polymerase-α/Primase (Pol-Prim) to initiate synthesis of RNA/DNA primers (20-22). Replication Factor C (RFC), Proliferation Cell Nuclear Antigen (PCNA), DNA polymerase delta, primer processing enzymes, and DNA Ligase I are recruited through interactions with the nucleic acid structures synthesized by LT/Pol-Prim/RPA/TopoI to synthesize the majority of the genome, remove RNA primers, and generate the covalently closed replicated episomes (22-27).

Much like cellular DNA replication, SV40 DNA replication is dramatically inhibited following treatment of cells with various types of DNA damaging agents, and this response is dependent on cellular DDR pathways. This inhibition can be observed on replicating SV40 DNA in living cells, as well as through decreased viral DNA replication in extracts from cells that had been subjected to DNA damage prior to preparation of cell extracts (13,14,28-37). Like inhibition of cellular DNA replication upon DDR, inhibition of SV40 DNA replication occurs through the action of trans-acting factors (12, 14, 17, 18) at both the initiation (14, 17) and likely the elongation (11,12,19) stages of replication (Fig. 1). This makes SV40 a valuable model system for investigating the mechanisms of human DDR and DNA replication checkpoints. Treatment of SV40 LT-expressing HEK293T cells with etoposide (ETO), a chemotherapeutic agent that inhibits cellular topoisomerase II and causes dsDNA breaks (DSBs), leads to dramatic inhibition of SV40 DNA replication through trans-acting mechanisms, and increased phosphorylation of LT by human DDR kinases ATR/ATM on one or more SQ/TQ sites (13). Conversely, viral DNA replication and helicase phosphorylation of a very similar viral DNA replication system, HPV16, which relies on many of the same host DNA replication proteins as SV40 (38,39), were both unaffected by ETO treatment (13). These findings suggested that ATR/ATM phosphorylation of LT could play a unique role in the inhibition of SV40 DNA replication. ATM and ATR, the apical regulatory kinases of the DDR, phosphorylate hundreds of effector proteins, preferentially on SQ or TQ amino acid (aa) motifs, to facilitate maintenance of genome integrity following DNA damage (40,41). ATM and ATR are also known to directly phosphorylate components of host DNA replication machinery, including the human replicative helicase proteins, MCM2-7, which play roles in cellular replication analogous to those of LT in SV40 DNA replication (42-45), in addition to the known effects of phosphorylation of CMG replication fork components by other checkpoint kinases (15,16). However, the functions of these DDR-dependent phosphorylation events on cellular replication proteins remain largely un-elucidated. Investigation of the effect of ATR/ATM phosphorylation of SV40 LT on viral DNA replication may provide insights into how ATR phosphorylation of the MCM proteins might affect DNA replication fork function in host cell DNA replication.

LT samples immunoprecipitated from LT-expressing human cells, both treated with ETO as well as mock-treated, were analyzed using tandem mass spectrometry (MS) to identify sites of LT phosphorylation. One of the SQ/TQ phosphorylation sites on LT occurred within its DNA helicase domain at threonine 518 (T518). Alignment of sixteen mammalian PyV LT aa sequences showed SV40 LT T518 to be highly conserved, being a TQ or SQ on all LT sequences aligned. We evaluated the DNA replication functions of phospho-T518 LT using a phosphomimetic aspartic acid mutant, T518D, in both cell-based and *in vitro* analyses. The vast majority of LT functions evaluated remain unaffected by this phosphomimetic mutation. However, the T518D phosphomimetic mutation dramatically inhibited the DNA helicase activity of LT, which would explain how SV40 DNA replication forks can be slowed following DDR.

## RESULTS

### Identification of novel SQ/TQ phosphorylation sites on SV40 Large T-antigen (LT)

Phosphorylation is known to modulate various functions of many DNA replication factors including SV40 LT. With regard to SV40 DNA replication, phosphorylation of threonine 124 (T124) on LT by the cell-cycle kinases, cdc2 or cdk2, is known to be important for LT’s binding to SV40 ori and assembly into a helicase (46-49); indeed this is a likely target for inhibition of the initiation of SV40 DNA replication through the well-established pathway from ATR to Chk1 to Cdc25 that inhibits cdc2/cdk2 activity (1,2,31,34,50). Phosphorylation of residues S106, S123, S639, S677, S679 and T701 have been observed, but appear to have little effect on LT’s DNA replication function (49); phosphorylation of S111/112 by CK2 or S120 by ATM have been shown to have some positive effect on SV40 DNA replication by increasing the efficiency of the adjacent LT nuclear localization sequence (NLS) (51-53). Since we previously showed DDR-enhanced phosphorylation of SQ/TQ residues on LT are associated with inhibition of SV40 DNA replication function (13) while the only previously demonstrated effects of SQ/TQ phosphorylation of LT were shown to have positive effects on SV40 DNA replication (51-53), we hypothesized that a hitherto unidentified DDR SQ/TQ phosphorylation site on LT might be mediating the DDR-inhibition of SV40 DNA replication. The SV40 LT protein sequence was submitted to NetPhos 3.1a which predicts the likelihood of in vivo aa residue phosphorylation by a large panel of well-studied human protein kinases based on aa sequence and protein structure (54). The results from the NetPhos prediction are shown in Supplemental Figure 1 with the S and T residues within SQ/TQ motifs clearly identified (Fig1). Note that all S and T residues within SQ/TQ motifs clearly identified (S120, T518, S639, S665, S667 and S677) in the six conserved SQ/TQ sites (residues S120, T518, S639, S665,S667 and S677) fall above the NetPhos Phosphorylation threshold Score of 0.500, indicating that all six are predicted likely to be phosphorylated in vivo (Fig 1).

Of the six SQ/TQ sites on LT that are all predicted by NetPhos to be phosphorylated, to date only three, S120, S639, and S677, have been demonstrated to be phosphorylated (32,49). To test the NetPhos prediction that all six SQ/TQ sites are phosphorylated in vivo we utilized a highly sensitive liquid chromatography-tandem mass spectroscopy (LC-MS/MS) approach to map phosphorylation sites on SV40 LT immunoprecipitated (IPed) from HEK293T cells. Because of the previously reported enhancement of LT SQ/TQ sites following DNA damage, LT was IPed from both mock- and etoposide (ETO)-treated HEK293T cells and immunoblotted using a commercial phospho-SQ/TQ preferential antibody (pSQ/TQ). Our results showed that slightly higher levels of LT were IPed from mock-treated cells, but that LT IPed from ETO-treated cells show higher levels of signal with the pSQ/TQ antibody (Fig 2A). These results were consistent with the antibody specifications that show substantial pSQ/TQ signal in normal cycling cells with enhanced reactivity following DDR. Figure 2B shows this enhancement of pSQ/TQ detection of global proteins in crude HEK293T extracts from treated cells (mock vs ETO, lanes 1 vs 3). Detection of the IPed LT using both anti pSQ/TQ and anti-LT antibodies is shown (Fig 2B & C). These results confirm the previously published DDR-enhanced SQ/TQ phosphorylation of LT (13). (Note that LT-associated proteins can be observed both by silver staining and pSQ/TQ immunoblotting, showing that some LT-associated proteins are SQ/TQ phosphorylated (Fig 1B and 1D)). Tandem MS (MS2) was used to evaluate LT samples from both mock- and ETO-treated HEK293T cells. LT was isolated by IP and subjected to trypsin digestion from either solvent (0.2% formic acid)-eluted IP pellets, or following SDS-PAGE and gel isolation. These analyses provided high coverage of the LT protein in all three samples analyzed (88.1% for OT, 91.4% for IT) with the identification of a large number of phosphorylated S/T/Y residues. Shown here are the identified pSQ/TQ sites identified on LT (Supl Fig 2). One example of a MS2 spectrum of one phospho-peptide is shown (Fig 1G), demonstrating phosphorylation of the LT T518 aa residue. Similar MS analyses of peptides from both IPed and gel-isolated LT detected phosphorylation at five of the six SQ/TQ sites, T518, S639, S665, S667 and S677 (Supl Fig 2, red boxes). The peptide containing S120, which was previously shown to be phosphorylated by ATM and to have a positive effect on nuclear translocation (32,53), was not detected by these MS analyses, so the phosphorylation of S120 could not be verified in this study.

**Fig 2.**
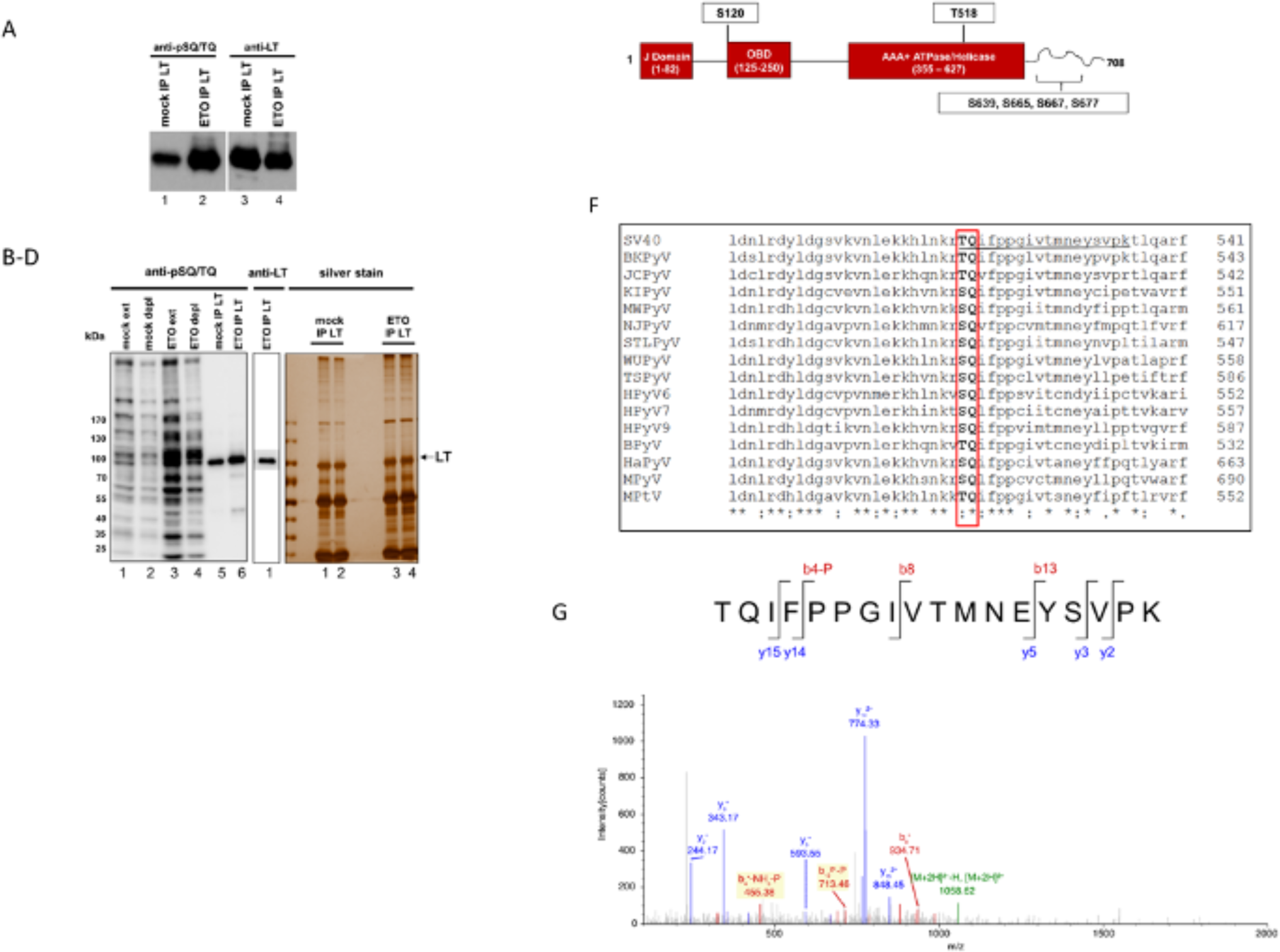
DDR-induced SQ/TQ phosphorylation sites on SV40 LT include the highly conserved T518. (A) Extracts from etoposide (ETO) treated or mock-treated HEK293T cells were subjected to immunoprecipitation of SV40 LT using monoclonal antibody Ab 419. SDS-PAGE was performed and immunoblotted with antibodies against LT (Ab 101) and a commercial antibody enriched for recognition of phospho-serine-glutamine and threonine-glutamine residues (anti-pSQ/TQ) as described in Methods. A 2-fold increase in pSQ/TQ signal (evaluated using Image J) was observed in the ETO lane compared to the mock lane when normalized for LT signal. (B) Immunoblot with anti-pSQ/TQ of extracts of mock and ETP-treated HEK293T cells, both prior and after LT immuno-depletion (lanes 1 and 2, 3 and 4), and of LT immunoprecipitations (as in panel A) (lanes 5 and 6). (C) Immunoblot with anti-LT (Ab 101) of the same sample as panel B lane 6, to verify SDS-PAGE migration of LT (note only the LT area of the gel is shown due to signal from the heavy and light chains of the antibody used in the IP). (D) protein stain (silver stain) of the samples in panel B lane 5 (lanes 1 & 2), and panel B lane 6 (lanes 3 & 4), which were excised from the gel for phospho-MS analysis. The un-labeled lane on the left shows the commercial molecular weight markers (indicated to the left of panel B). (E) A schematic of the 708aa SV40 LT sequence indicating the three major domains of LT (in red) and all SQ and TQ di-peptide sequences, the preferred DDR-induced ATM/ATR/DNA-PK substrate, that exist in the SV40 LT protein sequence. (F) LT aa sequences from 12 human polyomaviruses (SV40, BKPyV, JCPyV, KIPyV, MWPyV, NJPyV, STLPyV, WUPyV, TSPyV, HPyV6, HPyV7, and HPyV10), bovine polyomavirus (BPyV), hamster polyomavirus (HaPyV), murine polyomavirus (MPyV), and murine pneumotropic virus (MPtV) (indicated on the left) that surround the T518 aa residue of SV40 LT (top line). The highly conserved SQ/TQ di-peptide sequence is boxed in red, identical residues are indicated by the asterisk and conserved residues are indicated by a colon. The rightmost aa residue for each LT protein is indicated on the right. (G) One example of the dozens of MS/MS spectra demonstrating phosphorylated T518. This spectrum (Xcorr=1.76), of the peptide underlined in panel F, shows multiple product ions generated from the HCD fragmentation of the peptide backbone as shown. Notably, both b4 and b13 ions with neutral loss of the phosphate group have been detected (highlighted), indicating with high probability that the phosphorylated residue on this phosphorylated peptide is T518.

The large number of phosphorylation sites identified on LT (Supl Fig 2, lower panel) is consistent with the wealth of unexpected phospho-sites identified on many proteins using advanced mass spectrometric techniques in recent years (55,56). It has been estimated that only about a third of phospho-sites appear to have functional consequences (57). Our MS2 analyses showed five of the six of the SQ/TQ sites on LT to be phosphorylated in cells, and the remaining site had been previously shown to be phosphorylated and functional in enhancing LT nuclear translocation (32,49). It has been shown that evolutionary conservation of phospho-sites can be a useful approach to help identify which sites have functional significance (58). If arrest of DNA replication in response to DDR is important for SV40 and is mediated at least in part by phosphorylation of a specific SQ/TQ residue on LT, this aa residue is likely conserved in other PyV LT proteins. The Clustal Omega Multiple Sequence Alignment tool (The European Bioinformatics Institute (59)) was used to align sixteen mammalian PyV LT aa sequences, including: SV40, four other mammalian PyVs, and 11 human PyVs, focusing on conservation of the six SV40 LT SQ/TQ DDR kinase phosphorylation sites. These sites mapped to three distinct regions of the protein: to just upstream of the nuclear localization sequence (NLS) (S120) (Supl Fig3A), to within the C-terminal region (S639, S665, S667 and S677) (Supl Fig3B), and to the AAA+ ATPase/Helicase domain (T518) (Fig 2E). Our alignments revealed that the previously described S120 PIKK phosphorylation site in SV40 LT is highly conserved if one anchors alignment of this region to the predicted NLS of each of the LT sequences (see underlined aa residues, Supl Fig 3A). One or more SQ motifs exist four to ten aa residues upstream of all the NLS motifs (except for the human MWPyV which has two SQ motifs 24 and 28 aa residues upstream) (Supl Fig 3A). Hence the enhanced nuclear localization of LT due to phosphorylation of SV40 LT at S120 may be a common characteristic of the LTs of these other PyVs. The cluster of four SQ/TQ sites within the C terminus of SV40 LT is not well-conserved amongst these LTs. Many LTs, from both human and other mammalian PyVs, have few, or no SQ/TQ sites within this region (Supl Fig 3B). Studies of the SV40 LT C-terminal host range domain (aa 635-708) indicate that a C-terminal truncation of LT, to aa 635, are still viable for SV40 DNA replication (60). Together, these suggest that phosphorylation of one or more of the four C terminal SQ sites on SV40 LT would be unlikely to mediate inhibition of viral DNA replication, and may represent examples of the approximately two-thirds of phospho-sites that are phosphorylated without apparent functional consequences (57,60). Alignment of the PyV LT helicase domains showed that the SV40 LT phosphorylated TQ site, T518, is highly conserved, being an SQ or a TQ in all the helicase domains aligned (red box, Fig 2F). The MS2 peptide spectrum of one of the phospho-peptides containing T518, TQIFPPGIVTMNEYSVPK (underlined Fig2F), is shown (Fig 2G). This spectrum shows multiple product ions generated from HCD fragmentation of the peptide backbone. Notably, detection of both the b4 and b13 ions with neutral loss of the phosphate group indicate with very high probability that the phosphorylated residue on this phospho-peptide was T518. Forty events of T518 phosphorylation of this same peptide were detected, as well as dozens of events of phosphorylation of this site detected on larger peptides covering the T518 residue. With this LT phosphorylation data, the high degree of conservation of this PIKK phosphorylation site, the computational prediction by NetPhos, and the fact that T518 is the only SQ/TQ phosphorylation site within the enzymatic domain of SV40 LT, we hypothesized that phosphorylation of T518 on SV40 LT might affect LT’s ability to support viral DNA replication.

### Phosphomimetic mutant SV40 LT T518D is unable to support efficient viral DNA replication in human cells

To investigate whether phosphorylation at T518 could affect the ability of SV40 LT to support SV40 DNA replication, a phosphomimetic mutation at T518 was created by changing the threonine to an aspartic acid residue. This SV40 LT T518D mutation was evaluated for SV40 DNA replication function in human cells using a transient-transfection, luciferase-based SV40 viral DNA replication assay previously described by Fradet-Turcotte et al. (61). Briefly, human cervical cancer cells (C33a) are co-transfected with a SV40 LT expression plasmid (pCMV-sWT) and a SV40 ori-containing plasmid that also contains a firefly luciferase (Fluc) reporter gene (pFLORI40). Successful co-transfection results in expression of the LT protein which then binds to the SV40 ori on pFLORI40, driving replication of the plasmid. Higher levels of pFLORI40 template lead to higher levels of the Fluc reporter gene, resulting in higher expression of luciferase and higher Fluc activity. Mutant LT proteins compromised in SV40 DNA replication function result in lower levels of luciferase and lower Fluc activity. A third plasmid, pRL, drives expression of Renilla luciferase (Rluc); pRL does not contain the SV40 DNA replication origin. pRL is also co-transfected and serves as an internal standard for transfection efficiency and luciferase expression from a plasmid that is not replicated. SV40 DNA replication results in a dramatic increase in the ratio of Fluc to Rluc activities, and this Fluc:Rluc ratio has been shown to be a robust and sensitive assay for viral DNA replication (61). Plasmids expressing WT LT (pCMV-sWT) or a phosphomimetic mutant, T518D, (pCMV-sT518D) were evaluated thusly (Fig3A). We observed that phosphomimetic LT mutant pCMV-sT518D is severely deficient in supporting SV40 DNA replication (∼ 8 - 18% of WT LT DNA replication levels), while mutation of T518 to a non-phosphorylatable aa at that residue (alanine, T518A encoded by pCMV-sT518A) replicates at WT or slightly higher levels (Fig3A). Immunoblotting was used to verify that all three vectors expressed LT at comparable levels (Fig3B). These data show that an acidic substitution, but not a small uncharged substitution, at T518 inhibits the ability of LT to support efficient SV40 DNA replication in human cells, consistent with phosphorylation of this residue inhibiting SV40 DNA replication.

**Fig 3.**
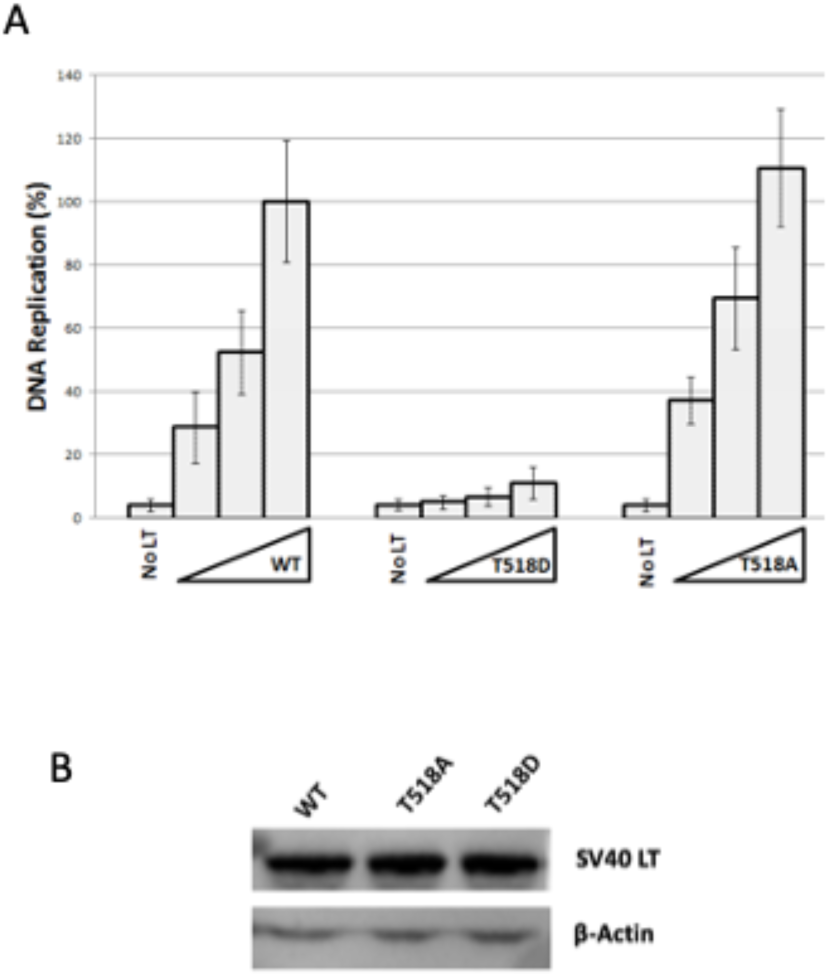
Mutant LT T518D is unable to support efficient SV40 DNA replication in living cells. (A) Increasing amounts (2.5, 5, and 10ng, as indicated by triangles on x axis) of pCMV-sWT, pCMV-sT518D, or pCMV-sT518A SV40 LT expression vectors were co-transfected into C33a cells along with a SV40-origin containing plasmid linked to firefly luciferase (pFLORI40) and a control plasmid lacking a viral origin linked to Renilla luciferase (pRL). After 72 hours, cells were harvested and assayed for the ratio of Firefly to Renilla luciferase to obtain relative SV40 DNA replication values for each triplicate LT construct well as described in the Methods and previously (28). No-LT controls show reliance of Firefly luciferase expression on LT. All values are reported as a percentage of the average values obtained with 10ng pCMV-sWT, which was tested in triplicate in each experiment. Results are an average of four separate experiments. (B) Western blot confirms comparable expression of SV40 WT, T518D, and T518A constructs in C33a cells. Error bars represent standard deviations.

### Phosphomimetic mutant SV40 LT T518D is fully functional as a transcriptional transactivator in human cells

To begin to determine whether the DNA replication deficiency of mutant LT T518D was specific, or due to an overall loss of structure/conformation, the ability of LT T518D to support LT’s transcriptional activation functions was investigated. The fact that the T518D mutant was expressed stably and not degraded was consistent with maintenance of the overall structure of LT T518D (Fig 3B). If LT T518D was capable of acting as a transcriptional activator this would verify that it was still capable of binding its dsDNA penta-nucleotide GAGGC binding sites within the SV40 origin/promoter control region (SV40 nt 5097-272), and the ability to act as a transcriptional activator would further confirm that LT T518D adopts a generally functional protein conformation. A vector was created that positions the SV40 late promoter (which requires LT binding for transcriptional activation) to drive expression of Renilla luciferase. The CMV promoter was removed from the pRL plasmid and replaced with nt 5097-272 of the SV40 genome from plasmid pSV010(-), so that the SV40 late promoter is oriented to control transcription of the Renilla luciferase gene. The resulting plasmid is referred to as pRLSV40-Late (Fig 4A). To circumvent an apparent decrease in transcription due to a loss in DNA replication function by LT T518D, this construct has a defective SV40 DNA replication origin due to a 4nt deletion in a critical LT binding site within LT binding domain II of the SV40 core origin which does not affect late promoter transcription (62). Human C33a cells were co-transfected with pRLSV40-Late along with increasing amounts of the WT or LT T518D expression vectors used in Fig 3. Cells were harvested after 72 hours and Renilla luciferase levels were evaluated to determine LT activation of SV40 late promoter transcription. Consistent with the known requirement of LT for SV40 late promoter activation, there was a substantial increase in transcription when increasing levels of the WT LT expression vector were co-transfected with the reporter construct (Fig 4B, WT). Increasing levels of the expression vector for T518D mutant LT resulted in essentially identical levels of transcriptional activation from the SV40 late promoter (Fig 4B, T518D). No appreciable difference in Renilla luciferase levels between WT and LT T518D in these assays indicates that the T518D mutant LT can still activate transcription from the SV40 late promoter, demonstrating that the T518D mutation does not adversely affect LT’s dsDNA binding or transcriptional activation functions.

**Fig 4.**
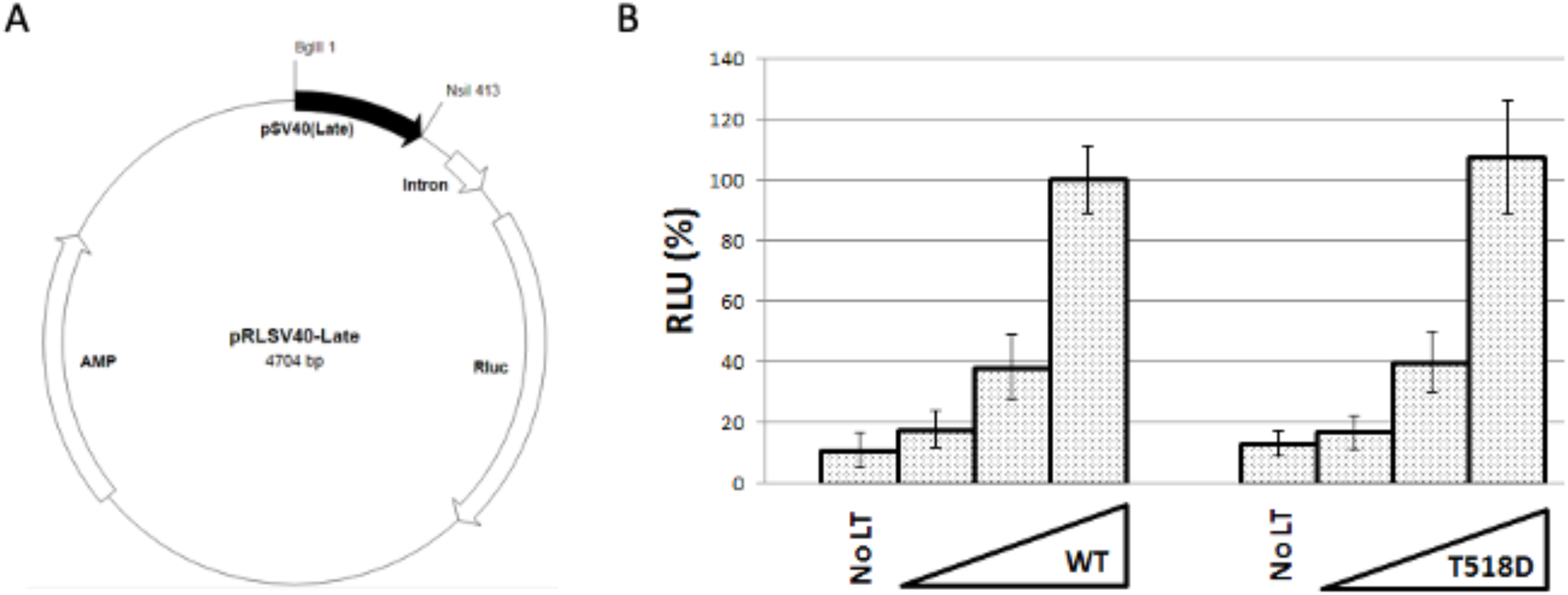
Mutant LT T518D is not deficient for binding to or activating transcription from the SV40 late promoter in living cells. (A) Plasmid map of pRLSV40-Late, highlighting SV40 origin/promoter sequence (SV40 genome nt 5097-272) oriented with the LT-dependent late promoter side driving expression of a Renilla luciferase gene. Note the small triangle within the SV40 sequence that indicates a 4-nt deletion that renders the SV40 origin inactive, allowing this construct to be used to monitor SV40 LT transcriptional activity in this cell-based assay without confounding effects due to SV40 DNA replication (27,30). Ampicillin resistance gene and an intron that increases splicing efficiency are also noted. (B) C33a cells were transfected with pRLSV40-Late, containing a Renilla luciferase gene driven by the late SV40 late promoter, and increasing amounts (25, 50, or 100ng) of expression plasmids for WT or T518D SV40 LT (as indicated by triangles on the x axis). Cells were harvested 72 hours post transfection and assayed for Renilla luciferase expression. No-LT controls demonstrate reliance of Rluc expression on presence of LT. All values are reported as a percentage of values obtained with 100ng pCMV-sWT expression vector, which was tested in triplicate in each experiment. Results are an average of three separate experiments, error bars represent standard deviation.

### Mutant SV40 LT T518D is unable to support efficient viral DNA replication *in vitro*

To evaluate the biochemical activities of LT required to support SV40 DNA replication and determine which function of LT is compromised by the T518D mutation, WT and LT T518D proteins were expressed and purified for biochemical analyses. LT requires phosphorylation at CDK site threonine 124 for SV40 DNA replication function. This phosphorylation does not occur if LT is produced in bacterial cells (63), therefore WT and LT T518D were expressed in and purified from High Five insect cells using baculovirus expression as described in the Methods (Fig 5A). The abilities of WT and LT T518D to support SV40 DNA replication *in vitro* are shown (Fig 5B). Increasing amounts of purified WT or LT T518D were combined with the SV40 origin-containing plasmid, pSV011, HEK293 human cell extracts (which contain all the other DNA replication proteins necessary to support SV40 DNA replication), and a buffer containing ribo- and deoxyribo-nucleotides and αP^32^dATP, for 90 minutes as described (Methods and (64)). Reaction products were subjected to neutral agarose gel electrophoresis and visualized using phosphorimager analysis. Addition of WT LT resulted in an increase in both fully replicated labeled plasmids (Fig 5B, Forms I, II and topoisomers, lanes 4-7), as well as in the episome multimers and the Cairns or “bubble structure” intermediates commonly seen with SV40 episomes replicated *in vitro* (Fig 5B, Replication Intermediates, lanes 4-7), as compared to control reactions containing either no LT (lane 3) or with the highest level of WT LT but with a plasmid template containing the mutated SV40 origin described above (lane 2) (62). Conversely, in parallel reactions with the same levels of LT T518D, synthesis of SV40 DNA replication products was severely deficient as compared to the WT LT (Fig 5B, lanes 8-11). To quantify the difference in SV40 DNA replication ability, reactions were spotted on DE-81 charged filter paper, washed, and incorporated P^32^dAMP was quantified by scintillation counting (as in the Methods). Consistent with the cell-based results, mutant LT T518D (open circles) was severely compromised in supporting SV40 DNA replication *in vitro*, showing maximum SV40 DNA replication levels of ∼20% of those synthesized with WT LT (open triangles) across a range of LT concentrations (Fig 5C). The control reaction using the SV40 replication origin mutant plasmid pSV011(-) at the highest level of WT LT, confirmed the dependence of this assay on a functional SV40 origin of replication (open square). These results were highly consistent with the cell-based SV40 DNA replication reactions, showing similar levels of synthesis by the phosphomimetic mutant LT as compared to WT LT.

**Fig 5.**
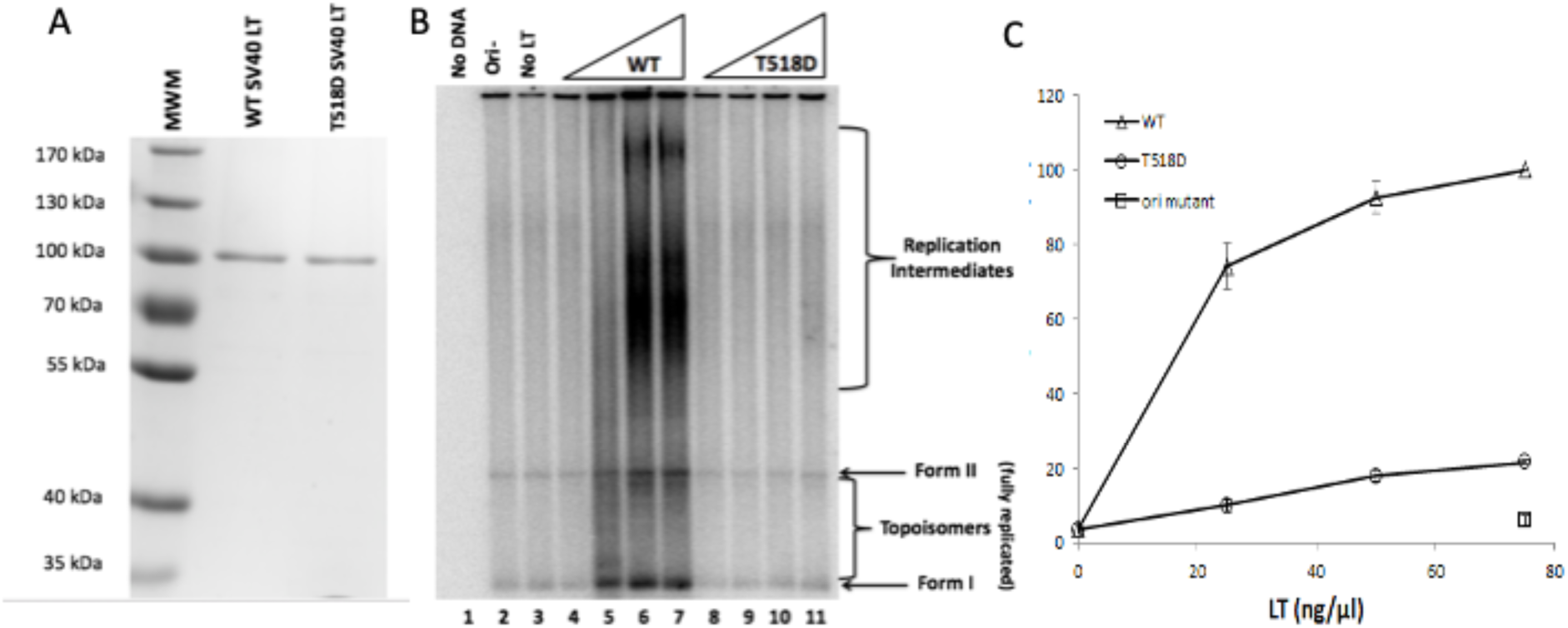
Mutant LT T518D is unable to support efficient SV40 DNA Replication *in vitro*. (A) Coomassie stain of WT and T518D SV40 LT expressed in and purified from insect cells through baculovirus expression, used for subsequent *in vitro* analyses in this study. (B) For *in vitro* DNA replication assays: increasing amounts (250, 500, 750, or 1000ng as indicated by increasing triangles) of purified WT or T518D LT were combined with the pSV011 SV40 origin-containing plasmid, HEK293 cell extract, and reaction buffer containing NTPs, rNTPs and αP^32^dATP. Reactions were resolved by agarose gel electrophoresis, followed by gel drying and exposure to phosphor-imager screens. No SV40 plasmid DNA was added in “No DNA” controls. pSV011(-) mutant SV40-origin plasmid was combined with the highest level of WT LT (the level used in lane 7) for “Ori-” control (lane 2). “No LT” control (lane 3) shows background incorporation of αP^32^-dATP into pSV011 SV40-origin plasmid by host DNA repair machinery in the absence of LT. Replication intermediates signify ongoing DNA replication. Form I/II DNA and topoisomers signify fully replicated plasmid DNA, and replication intermediates indicate Cairns intermediates and interlocked not-yet-resolved episomes. (C) Incorporation of P^32^dAMP into newly synthesized SV40 DNA was quantified by spotting of reactions on DE81 filter paper and scintillation counting. All values are reported as percentage of the value obtained with 75 ng/μl of WT LT in that experiment. Results are an average of three independent experiments. Ori- (square) is a negative control reaction using the highest amount of WT LT with SV40 origin-mutant plasmid pSV011(-).

### Mutant T518D interacts with host cellular replication factors and stimulates Pol-Prim polymerase activity like WT LT

To initiate SV40 DNA replication, LT must bind ATP, oligomerize into hexamers, and make crucial contacts with host RPA and TopoI, in addition to binding to, and stimulating the DNA polymerase activity of, host Pol-Prim (20,65-68). To address the ability of phosphomimetic LT to interact with cellular DNA replication factors, binding of WT and LT T518D to RPA, TopoI, and Pol-Prim was evaluated using enzyme-linked immunosorbent assays (ELISA) (Fig 6). Human TopoI (Fig 6A) or RPA (Fig 6B) were immobilized in 96 well plates; after blocking, wells were incubated with increasing amounts of WT LT (open triangles) or LT T518D (open circles), and binding was detected with monoclonal antibody against LT and HrP-linked secondary antibody. WT and T518D interacted with both human TopoI and RPA to the same degree (Fig 6A & B). Controls show that binding of LT was clearly dependent on TopoI and RPA, respectively (Fig 6A & B, open squares). Pol-Prim ELISAs were carried out by immobilizing constant levels of either WT LT or LT T518D to the wells, and after blocking, incubating with increasing amounts of Pol-Prim, which was then detected using a Pol-Prim primary antibody and HrP-linked secondary antibody (Fig 6C). As observed with human TopoI and RPA, we observed no difference between WT and LT T518D for Pol-Prim binding (open triangle versus open circle), and binding of Pol-Prim was clearly dependent on immobilized LT (open square) (Fig 6C). Since LT’s interaction with p53 also occurs within the LT helicase domain, we also evaluated whether the interaction with p53 was affected by mutation at T518. LT, as well as the T518 point mutant proteins were expressed in human cells, IPed and immunoblotted for LT and p53. The LT proteins were precipitated to a similar degree and were equally capable of co-IPing cellular p53 (Supl Fig4).

**Fig 6.**
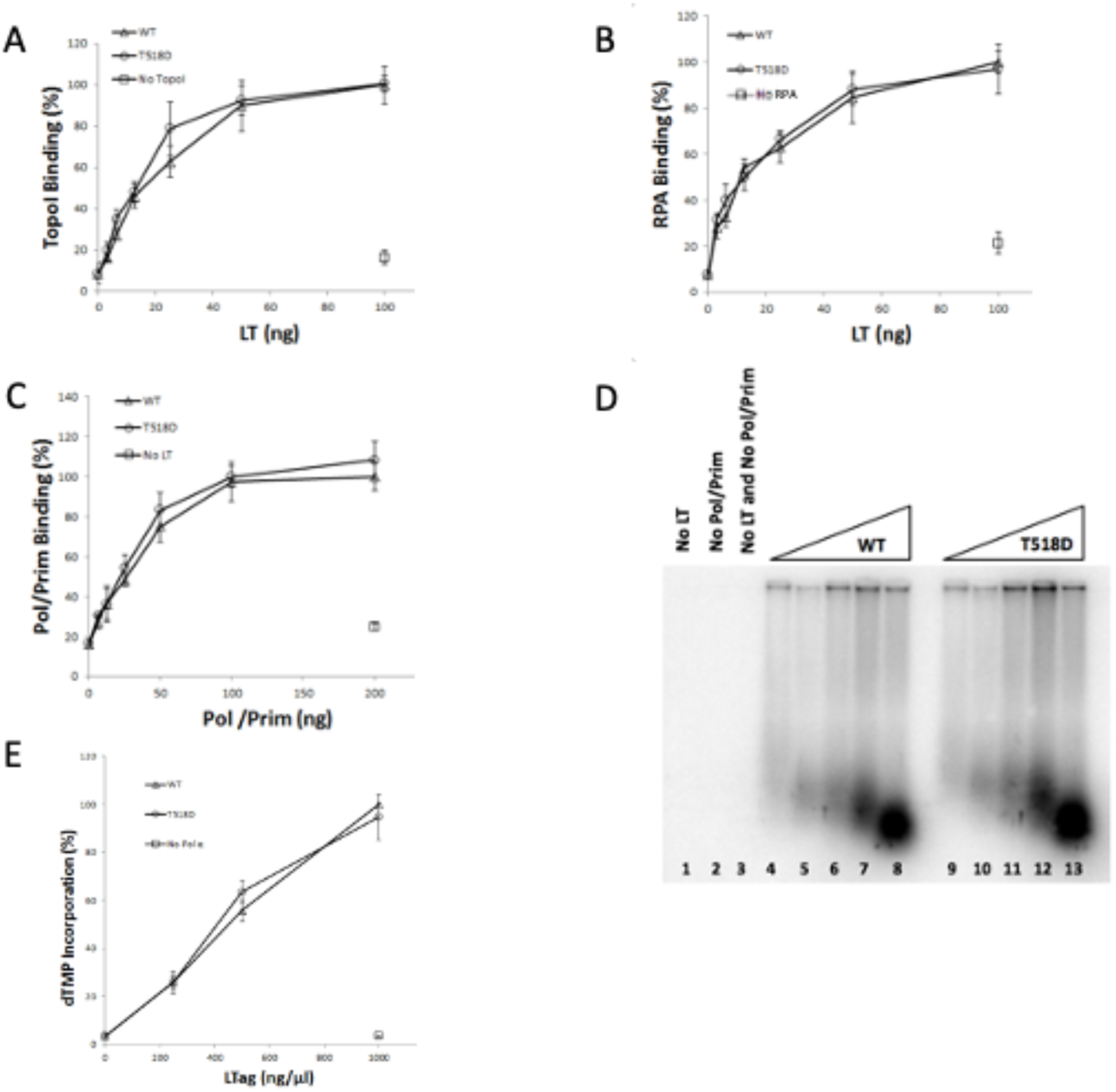
The SV40 LT T518D mutant protein is proficient for binding and stimulating host DNA replication proteins. (A) TopoI or (B) RPA were immobilized in 96 well plates followed by addition of increasing amounts (3.125, 6.25, 12.5, 25, 50, or 100ng) of WT (open triangles) or T518D (open circles) LT as a secondary protein as indicated by the X axes. ELISAs were completed with a primary pAB101 LT antibody followed by a secondary HrP antibody. Block control (open square) represents non-specific binding of WT LT to blocking protein mixture (5% milk and 2% FBS in TBST). All values are reported as a percentage of values obtained from the interaction of 100ng WT LT with immobilized TopoI (A) or RPA (B). (C) For Pol/Prim ELISAs: WT or LT T518D were immobilized in 96 well plates followed by addition of increasing amounts (6.25, 12.5, 25, 50, 100, or 200ng) of human Pol/Prim as a secondary protein as indicated by the X axis. ELISAs were completed with a primary mAB1645 Pol/Prim antibody followed by a secondary HrP antibody. Block (open square) represents non-specific binding of Pol/Prim to blocking protein mixture. For panel C, all values are reported as a percentage of values obtained from the interaction of 200ng Pol/Prim with immobilized WT LT. For panels A-C, each ELISA shown is a representative experiment with each binding reaction run in triplicate. (D) For Pol/Prim stimulation assays: reactions were assembled with increasing amounts (62.5, 125, 250, 500, or 1000ng) of WT LT (lanes 4-8) or LT T518D (lanes 9-13) as indicated by triangles, a constant amount of Pol/Prim, poly[dA]/oligo[dT] substrate, and αP^32^dTTP. Reactions were resolved by denaturing agarose gel electrophoresis. Gels were dried and exposed to phorphor-imaging screens. Lane 1 is a control lacking LT but containing 50ng Pol/Prim, showing the heavy reliance of the assay on the presence of LT at these low levels of Pol/Prim. Lane 2 is a control lacking Pol/Prim but containing 1000ng of WT LT, showing the reliance of the assay on Pol/Prim. Lane 3 is a control lacking both LT and Pol/Prim. (E) Stimulation of Pol/Prim by LT was quantified by scintillation counting of reactions spotted on DE-81 filter paper, washing of unincorporated nucleotides, and scintillation counting as above. All values are reported as percentage of the value obtained with 1000ng WT LT in that experiment. Results are an average of three separate experiments, the error bars represent standard deviation.

Because the DNA polymerase activity of Pol-Prim is stimulated by the interaction with LT, a DNA polymerase assay using a poly dA/oligo dT template was used to determine whether T518D could stimulate Pol-Prim (20,22). Pol-Prim was combined with increasing amounts of WT LT (lanes 4-8) or LT T518D (lanes 9-13) with a poly dA/oligo dT template and αP^32^dTTP (Fig 6D). Reactions were resolved on denaturing gels, exposed to phosphorimaging screens, and incorporation of P^32^dTMP into DNA polymers by Pol-Prim was analyzed by phosphorimage analysis. Controls (Lanes 1-3) show that appreciable DNA synthesis is reliant on the presence of both SV40 LT and minimal levels of human Pol-Prim (extremely low levels of Pol-Prim were used in these assay conditions to maximize sensitivity of the LT stimulation). LT T518D was capable of stimulating DNA polymerase activity of human Pol-Prim equally as well as WT LT, shown by equivalent increases in ssDNA products containing incorporated P^32^dTMP detected via denaturing gel electrophoresis. Quantification of three separate Pol-Prim stimulation assays is shown (Fig 6E). As SV40 LT is only known to interact with human replication factors, TopoI, RPA, and Pol-Prim, the results presented in Figure 6 suggest that the observed inability of mutant LT T518D to support SV40 DNA replication does not appear to be due to an inability to interact functionally with human DNA replication proteins.

### LT T518D is a functional ATPase, but is unable to efficiently unwind long dsDNA

The ATP-dependent DNA helicase activity of hexameric LT is essential for SV40 LT viral genome replication, and since T518 is located in the ATPase/helicase domain, both ATPase and DNA helicase functions of LT T518D were investigated. For ATPase assays, purified WT (open triangles) or T518D (open circles) LT hydrolysis of γP^32^ATP was assayed over time (Fig 7A), in the presence of increasing amounts of ssDNA (Fig 7B), and over a range of ATP concentrations (Fig 7C). Reactions were developed on TLC plates, exposed to phosphor-imaging screens, and quantified as described in the Methods. We observed that WT LT and LT T518D are both able to hydrolyze γP^32^ATP at the same rate over a 90 min time course as seen by equivalent increases in free, hydrolyzed γPO_4_^3-^ in both WT and T518D reactions (Fig 7A, representative TLC plate phosphorimage shown in inset). The ATPase function of LT is known to be stimulated by binding to ssDNA, so the ability of WT and LT T518D to hydrolyze ATP was examined in the presence of increasing amounts of polydT (65). WT and T518D ATPase function were stimulated to equivalent degrees by addition of varying amounts of polydT (Fig7B). This suggested both proteins interact similarly with ssDNA. To evaluate the ATPase activities of both WT and LT T518D at varying levels of ATP, ATP hydrolysis was evaluated in the presence of increasing concentrations of unlabeled ATP. As levels of unlabeled ATP were increased in these reactions, we observed an apparent decrease in the amount of γP^32^ATP hydrolyzed (Fig7C) (note that the overall levels of ATP hydrolyzed actually increased in these reactions because more hydrolysis of unlabeled ATP is occurring). In these reactions we observed equivalent decreases in the proportion of γP^32^ATP hydrolyzed in both WT and LT T518D reactions, demonstrating that WT and LT T518D showed equivalent ATPase activity over a range of ATP concentrations, indicating that the two LTs have comparable affinities for ATP. Figure 7 demonstrates that despite mutant LT T518D’s inability to support efficient SV40 DNA replication, LT T518D retains WT ATPase activity over a range of conditions.

**Fig 7.**
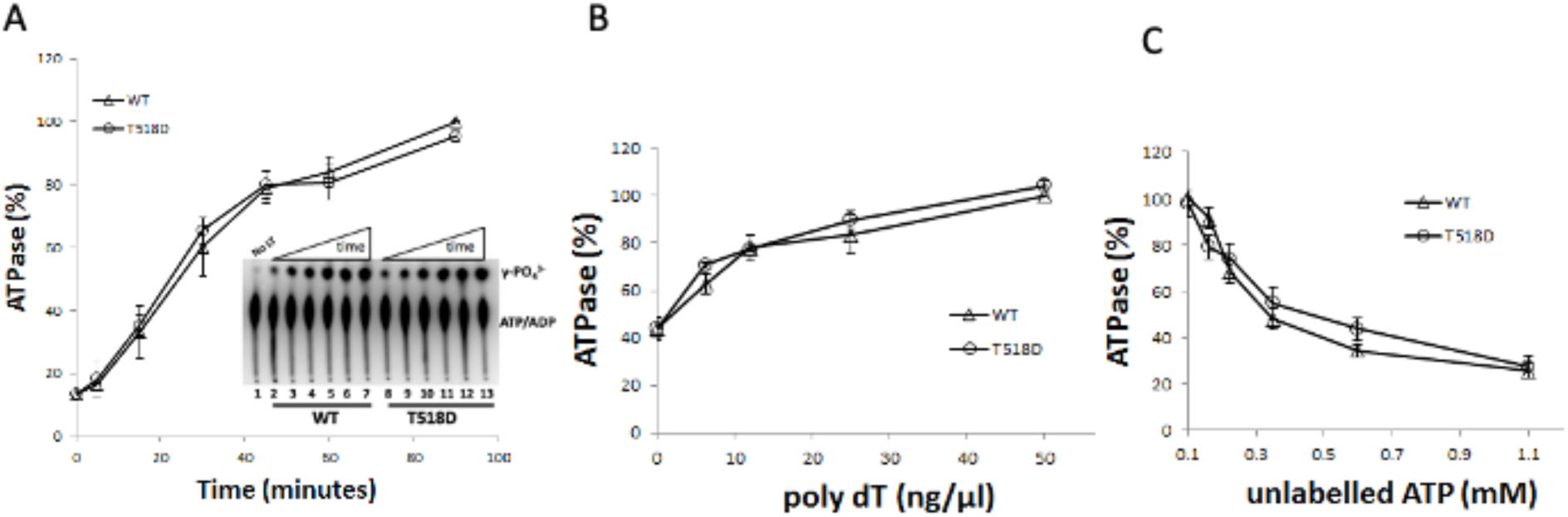
LT mutant T518D is proficient for ATPase function *in vitro*. (A) WT or LT T518D was incubated with γP^32^-ATP at different points (5, 10, 15, 30, 45, 60 and 90 minutes) over a time course (as indicated by increasing triangles). Reactions were subjected to TLC and plates were exposed to phospho-imaging screens, followed by autoradiographic analysis. Representative ATPase time course TLC plate image is shown in inset. Lane 1 shows a control reaction without LT. Lanes 2-7 show WT LT ATPase reactions terminated at increasing time points over the course of 90 minutes, and lanes 8-13 show LT T518D ATPase reactions over the same time course. TLC images from three separate experiments were quantified using Image J software, averaged, and plotted above. Values are reported as a percentage of the density values of hydrolyzed γ-phosphate spots on TLC plates obtained with WT LT at the 90 minute time point. Results are an average of three separate experiments (B) WT (open triangle) or LT T518D (open circle) was incubated with γP^32^-ATP and an increasing concentration (0, 6.25, 12.5, 25, and 50ng/μl) of 221bp polydT single stranded DNA as indicated by the x axis. All values are reported as a percentage of the density values of hydrolyzed γ-PO_4_^3-^ spots on TLC plates obtained with WT LT at the highest poly dT concentration. Results are an average of three separate experiments. (C) WT or LT T518D was incubated with γP^32^-ATP in ATPase reactions with increasing concentrations (1, 0.6, 0.35, 0.225, 0.1625, or 0.1mM) of unlabeled ATP as indicated by the x axis. All values are reported as a percentage of the density values of hydrolyzed γ-phosphate spots on TLC plates obtained with WT LT at 0.1mM cold ATP reaction concentration. Results are an average of three separate experiments. Error bars represent standard deviations.

Hydrolysis of ATP leads to conformational changes within DNA-bound hexamers of LT, leading to translocation of LT along the leading strand template and separation of dsDNA during DNA replication (69). Therefore, we assessed the DNA helicase activities of WT and LT T518D over time using two substrates with differing lengths of dsDNA: one composed of a short (17bp) duplex region (Fig8A) and another with a longer (100bp) duplex region (Figs 8B & C). dsDNA substrates separated in helicase reactions were resolved on native TBE polyacrylamide gels and subjected to phosphorimage analysis. In each gel image, control lane 1 (no LT) shows migration of the fully duplex DNA substrate while lane 2 (boiled substrate) shows the migration of the radiolabeled strand of the substrate removed during the reaction. Over time, both WT (lanes 3-8 Fig 8A & 8B) and LT T518D (lanes 9-14 Fig 8A & 8B) unwind a short 17nt duplex region of DNA with equal efficiencies as seen by equivalent accumulation of released 17nt oligo. However, LT T518D is much less efficient than WT LT when the region of duplex DNA is increased to 100bp. Figure 8B is a representative gel showing that over time less 100bp helicase substrate is unwound by T518D than by WT LT. Figure 8C is a quantification of three separate experiments showing that LT T518D unwinds the 100mer substrate with only 35%-40% of the efficiency of WT LT over time. This difference appears to be more dependent on the difference in length of the DNA being unwound rather than the structure of the template, as studies with a forked substrate, like the 100 base pair substrate, but with only 50 paired bases, showed an intermediate result between results with the 17bp and the 100bp substrates (Supl Fig5, rather than varying time, these results show varying levels of LT, WT and T518D, added to helicase assays with 50bp or 100bp dsDNA forked substrates). LT T518D was more efficient at unwinding the shorter 50bp substrate than the 100bp substrate but was deficient compared to WT on both substrates. These data indicate that LT T518D is less efficient than WT LT at separating longer stretches of duplex DNA, while it is proficient at removing short segments of DNA.

**Fig 8.**
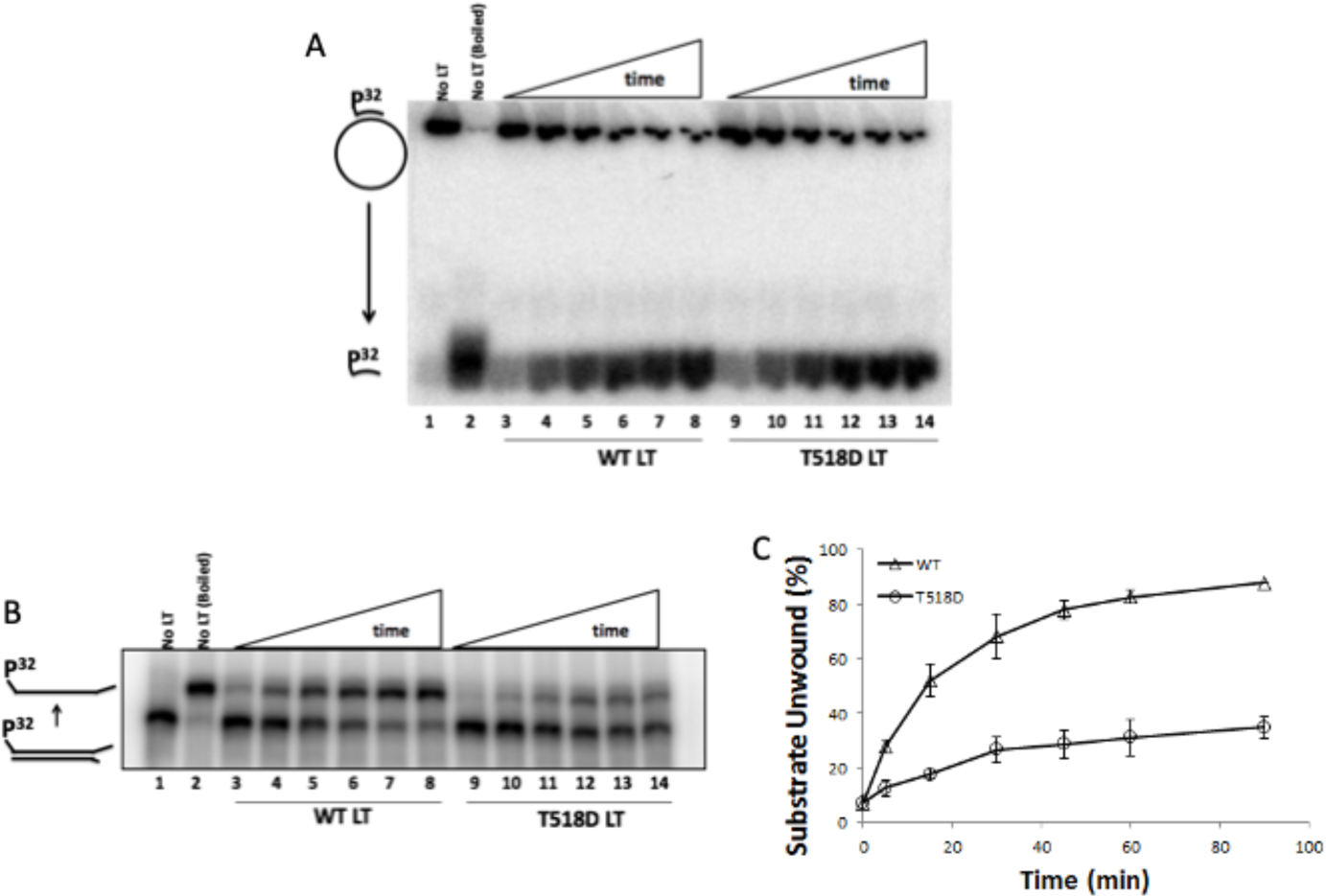
LT mutant T518D is deficient for progressive DNA helicase activity. WT or LT T518D were incubated with a dsDNA helicase substrate with (A) 17bp or (B) 100bp of duplex DNA, each having a single P^32^ end-labelled strand, at different time points (5, 10, 15, 30, 45, 60, or 90 minutes) over a time course as indicated by increasing triangles. Reactions were resolved on native TBE gels, exposed to phorphor imaging-screens, and analyzed by autoradiography. Values were obtained by setting the density of the released, radiolabelled band of a boiled reaction to 100% helicase activity and comparing densities of released bands in reactions to the boiled control. (A) and (B) are representative images for 17mer and 100mer time course helicase assays. “No LT” controls (lanes 1) show migration of intact, duplex DNA helicase substrates compared to “No LT (Boiled)” controls (lanes 2) showing migration of the single, radio-labelled strand of the helicase substrate following separation of DNA strands (note substrate cartoons highlighting the difference in direction of migration of the boiled 17mer and 100mer substrates). (C) represents quantification of three separate 100mer helicase time course assays. Error bars represent standard deviations.

### Phosphomimetic mutation of JCPyV LT is also compromised for viral DNA Replication

Our decision to evaluate the potential function of phosphorylation at the T518 residue of SV40 LT was based in part on the conservation of this site amongst PyV LTs (Fig2F); if DDR regulation of PyV DNA replication by phosphorylation at this site is important, then phosphomimetic mutations at this site in other PyV LTs should also show the phenotype of decreased viral DNA replication. A vector for the expression of LT from the clinically relevant human PyV, JCPyV, is available and was obtained and mutagenized to a phosphomimetic residue at this highly conserved site, JCV LT T519D. A dual luciferase human cell-based JCPyV DNA replication system was used to evaluate the relative ability of WT and T519D JCPyV LT proteins to support JC PyV DNA replication (70). Just as with SV40, mutation of JC PyV LT to a phosphomimetic residue resulted in levels of JC PyV DNA replication far lower than levels support by WT JC LT (Fig9).

**Fig 9.**
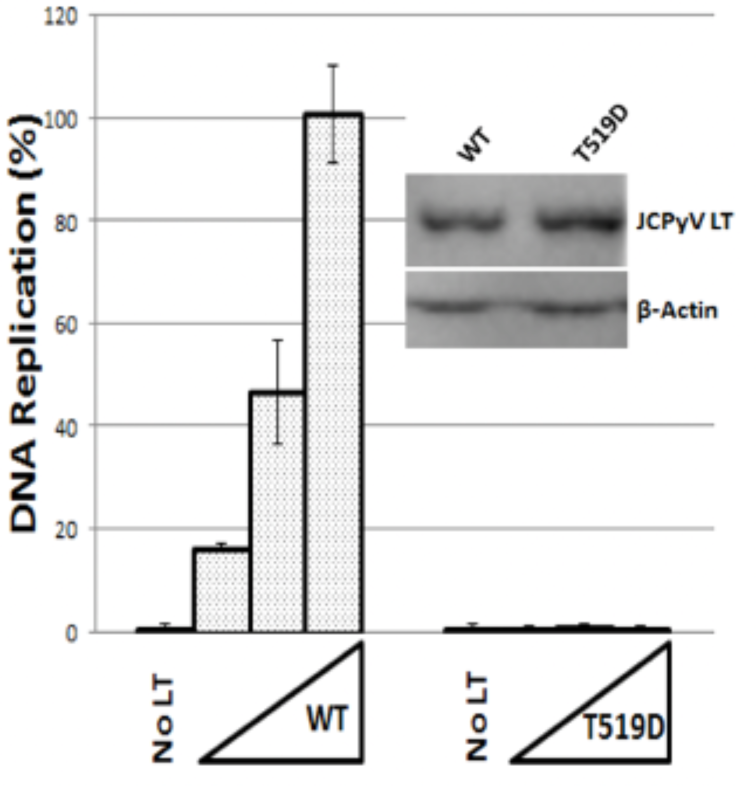
Phosphomimetic mutation of JCPyV LT, T519D, is deficient for JCPyV DNA Replication in human cells. Increasing amounts (2.5, 5, or 10 ng as indicated by X axis) of pCMV-jWT, or pCMV-jT518D JCPyV LT expression plasmids were transfected into C33a cells in 96 well plates with respective JCPyV origin-containing Firefly luciferase plasmid and a control plasmid lacking a viral origin linked to Renilla luciferase (pRL). After 72 hours, cells were harvested and assayed for the ratio of Firefly to Renilla luciferase to obtain relative JCPyV DNA replication values for each replicate well. No LT controls show reliance of Firefly luciferase expression on LT. All values are reported as a percentage of the values obtained with 10ng pCMV-jWT LT. Results are a single experiment, run in triplicate, representative of data from three separate experiments. Western blots confirm equal expression of each JCPyV LT construct in the same C33a cells used for the JCPyV experiments. Error bars represent standard deviations.

## Discussion

The SV40 DNA replication system is an important tool for studying the effects of DNA damage on both viral and eukaryotic DNA replication (13,14,28-32,34-37,71). Although strong activation of DDR may inhibit/pause or slow DNA replication, faithful duplication of both SV40 and human DNA is dependent on functional ATM and ATR kinases. ATM acts to prevent unidirectional replication and to alleviate concatemerization (72), while ATR acts to prevent origin firing in the presence of DDR, and as a surveyor to slow and/or stabilize moving DNA replication forks that have encountered a lesion, preventing fork collisions and DNA breaks (1,2,4,73,74). SV40 LT and human MCM helicase proteins are both phosphorylated by ATM/ATR following DNA damage, although the effect of most of these phosphorylation events remain unclear (13,32,42-45). Since these helicases are the central motor proteins of DNA replication forks, it is likely that DDR-dependent phosphorylation of helicases contributes to DDR-dependent inhibition of DNA replication by slowing or transient pausing of DNA replication forks to allow time to correct potentially lethal (or mutagenic) damage to DNA. Our aim was to study the DDR-dependent phosphorylation of a replicative DNA helicase and the impact of these phosphorylation events on DNA replication, using SV40 LT as a model helicase. MS2 was used to identify DDR-dependent SQ/TQ phosphorylation sites on SV40 LT. We show here for the first time that the T518 aa residue on SV40 LT is indeed phosphorylated in living cells (Fig2). The conservation of this aa residue on a variety of PyV LT proteins, and its presence within LT DNA helicase domain (Fig2), suggested an important role for this residue in DDR.

We investigated the likely effect of phosphorylation of the T518 residue of SV40 LT using a phosphomimetic mutation of LT, T518D. This mutation resulted in stable expression in human cells, did not show signs of any apparent increase in degradation, and retained wild-type function in cell-based LT-dependent transcriptional activation (Figs 3&4), all indicating that LT T518D is stably expressed and properly folded when expressed in human cells. Purified recombinant LT T518D also retained WT levels of binding to: TopoI (reported to contact LT at aa 83-160 and 602-708), Pol-Prim (reported to contact LT between aa 260-627), and RPA (reported to contact the LT OBD), further supporting the notion that LT T518D is being properly folded in both settings: expressed in human cells as well as expressed in insect cells and purified (Fig6) (75,76). The p68 “regulatory subunit” of Pol-Prim is known to interact with LT within its ATPase/helicase domain. Hence, we hypothesized that phosphorylation of LT T518 might disrupt the ability of LT to bind to and stimulate the DNA synthesis activity of Pol-Prim (77). However, LT T518D was able to bind and stimulate the activity of human Pol-Prim as well as WT LT in our ELISA and Pol-Prim DNA synthesis stimulation assay analyses (Fig6). The surface exposure and location of T518 near the interface of adjacent monomers in the SV40 LT hexamer structure initially suggested that mutation or phosphorylation at T518 might disrupt the ability of LT to oligomerize into functional hexamers, a crucial late step in initiation of DNA replication, as well as required for helicase function/progression. Since T518 is distal to the LT ATPase site (∼20 angstroms), mutation at this site might not be expected to affect ATPase function; however, decreased ability to hexamerize would compromise LT’s ATPase function. Hexamerization is important for ATPase/helicase function because the LT ATP binding/hydrolysis pockets are created by components of two adjacent LT monomers in the LT hexamer (78,79). However, our data showing that LT T518D is as active as WT LT in ATPase function (Fig7) indicates that LT T518D must also be proficient for hexamerization (this was consistent with proficient LT hexamerization by LT T518D on native PAGE, data not shown). In retrospect, since SV40 LT hexamerization is primarily dependent on contacts upstream of T518, in the LT origin-binding (OBD) (aa 125-250) and oligomerization domains, and is dependent on CDK2 phosphorylation of T124, perhaps this result should not have been surprising (46,80).

During DNA replication fork progression, it is proposed that SV40 LT hexamers surround the leading strand template bound to six ATP molecules in a “ready” state. Each ATP binding pocket consists of cis and trans elements belonging to adjacent LT subunits; a cis Walker A P-loop ATP-binding element from one monomer and the Walker B, arginine/lysine finger ATP hydrolysis elements from the adjacent monomer (69,79). In an ATP bound state, each LT subunit makes a unique base interaction with DNA through β-hairpins (aa 508-517 from each monomer) inside the hexamer’s central channel (78). Upon hydrolysis of ATP, conformational changes in LT near the β-hairpins cause translocation of the hexamer in the 3’ to 5’ direction along the leading strand template ssDNA. Further conformational changes occur between the ADP-bound and the second round of ATP-bound state of LT hexamers, causing further translocation of the hexamer and physical exclusion of the lagging strand template, which results in “helicase” displacement of the lagging strand template (69,78,79). As noted above, T518 is distal from the ATP binding pocket located at the interface between the LT monomers of the LT hexamer (∼20 angstroms). However, T518 is adjacent to the β-hairpin itself. This could explain how mutation/phosphorylation at this residue could compromise DNA helicase function without affecting ATPase function. Loss of proper β-hairpin function would not affect the ATPase-dependent conformational changes in the LT monomers but could compromise the mechanical connection between the ATP hydrolysis/LT conformational changes and the mechanical action required to track along the ssDNA. Since LT T518D is still capable of hexamerizing and hydrolyzing ATP (as well as binding to ssDNA, as evidenced by its wt levels of ssDNA stimulation of ATPase function), it should still be able to displace short stretches of ssDNA even without its motive force. Indeed, LT T518D was fully capable of removing a short (17 base) oligonucleotide annealed to a longer ssDNA template (Fig8A). However, lack of wild-type β-hairpin function/motive force of the helicase would not allow for efficient displacement of longer stretches of ssDNA from a longer dsDNA template. This is what was seen with oligonucleotides of 50 and 100 bases (Figs 8B, 8C, Supl5). Hence, it appears that mutation/phosphorylation at T518 may affect the ability of LT to coordinate ATP hydrolysis with the mechanical motion required of the DNA-interacting β-hairpin for helicase function. This conserved site is either a serine or a threonine in many PyV LTs, suggesting the length of the aa side change is not as critical as the change in charge caused by mutation/phosphorylation. The similarity in charge/general size is consistent with phosphorylated T518 displaying a similar mechanical defect in helicase progression as mutation to an acidic residue.

The fine details of the mechanisms of how phosphorylation of LT at T518 (or mutation to an aspartic acid) might result in compromising β-hairpin function or helicase slowing/pausing remain to be determined. One could imagine that compromising β-hairpin function could cause slippage during helicase progression as one or more LT monomers become modified, resulting in slowing or stalling of the replication fork, consistent with what is seen with DNA replication forks in mammalian cells following DDR activation (57, 58). Although this SV40 T518 site has no direct analogous SQ/TQ residue adjacent to the β-hairpins in the human MCM2-7 replicative helicase proteins, alignments of all six individual MCM’s 2-7 from humans, several vertebrates, and through fission and budding yeast reveal a highly conserved and uncharacterized SQ site located just downstream of the Walker A ATP-binding motifs of each protein. It is possible that the observed ATR/ATM phosphorylation of MCMs causes slowing or pausing of MCM helicase function, similar to what we have seen with SV40 LT T518D, and that could cause the well-established ∼10-fold decrease in replication fork rate seen when eukaryotic cells are subjected to DDR (7,81). It will be interesting to investigate the effect of mutations at DDR-dependent SQ/TQ phosphorylation sites on other recombinant replicative helicases including other PyV LTs and MCMs.

Human RPA is also phosphorylated following DNA damage, and it has been shown that RPA purified from cells subjected to DNA damage is deficient in its ability to support SV40 DNA replication *in vitro* when using naïve LT (29,34,71,82,83). Hence, we would not expect the LT T518A mutation to render SV40 DNA replication resistant to DDR-dependent slowing, as the DDR effect on RPA function would be sufficient to inhibit SV40 replication. It is not surprising that there would be multiple redundant pathways to slow DNA replication forks in response to DDR as redundancy is common in the cellular DDR response. Currently little is known about how DDR-dependent phosphorylation of RPA renders it deficient for SV40 DNA replication. Since RPA interacts with SV40 LT and cooperates with LT during helicase function (84), it is possible that phosphorylation of RPA acts through LT helicase function. Conversely, RPA is critical for priming and DNA synthesis through its interactions with LT and Pol-prim (20-22), so it is possible that this is the function modulated by RPA phosphorylation. It is attractive to propose that inhibition of the elongating DNA replication fork occurs through two independent mechanisms, one on the leading strand (through LT helicase inhibition) and one on the lagging strand (by prevention of primer synthesis through phosphorylation of RPA inhibiting required protein-protein interactions). Having redundant pathways for DDR targets is not unusual (1,2,4,74), and having pathways that that work on both leading and lagging strand processes would likely lead to more stably paused replication forks. Based on these results as well as previous studies it is likely that phosphorylation of LT or host DNA replication factors (such as RPA) may be capable of suppressing SV40 DNA replication individually, and that two or more redundant pathways have evolved to slow or pause DNA replication following activation of DDR.

## METHODS

### Plasmids, Mutagenesis, DNA substrates

WT LT-encoding baculovirus transfer vector p941-WT was a gift from Dr. Dan Simmons (University of Delaware). Mutant LT T518D-encoding transfer vector p941-T518D was created by site-directed mutagenesis using the primers: 5’-GAAACACCTAAATAAAAGAGATCAAATATTTCCCCCTGGAATAGTCACC-3’ and 5’-GGTGACTATTCCAGGGGGAAATATTTGATCTCTTTTATTTAGGTGTTTC-3’ (Integrated DNA Technologies), and verified by DNA sequencing. An aspartic acid was used instead of a glutamic acid as the phosphomimetic residue for threonine because many LTs in our alignment (Fig 2F) showed a serine at this site rather than a threonine, so we surmised that the charge change upon phosphorylation was the more important aspect than the side chain length, and using a slightly shorter side-chain length would be less likely to create unanticipated steric issues due to the differences in bond angles between the atoms of the glutamic acid and phospho-threonine side chains.

pSV011 contains an SV40 origin fragment (nucleotides 5171-128) in pUC18 (85). pSV011(-) is the same SV40 origin plasmid with a 4nt deletion in LT binding site I (SV40 nt 5239-5242), rendering this plasmid deficient for DNA replication (62).

SV40 origin-containing Firefly luciferase plasmid (pFLORI40) and pCMV expression vector for WT SV40 LT (pCMV-sWT) as well as a Renilla luciferase plasmid (pRL)) were gifts from Dr. Amelie Fradet-Turcotte (University of Laval) and Jacques Archambault (McGill University) (61). JCPyV origin-containing Firefly luciferase plasmid (pFLORIjc) and pCMV expression vector for WT JCPyV LT (pCMV-jWT) were also provided as gifts from Dr. Jacques Archambault (McGill University) and Dr. Peter Bullock (Tufts University) (70).

Mutant pCMV-sT518D was created by site directed mutagenesis with a pCMV-sWT template using the same primers used to generate p941-T518D. Mutant SV40 and JCPyV pCMV plasmids were created using site directed mutagenesis with respective WT plasmids for templates using the following primer pairs: pCMV-sT518A (5’-GAAACACCTAAATAAAAGAGCTCAAATAATTCCCCCTGGAATAGTCACC-3’ and 5’-GGTGACTATTCCAGGGGGAAATATTTGAGCTCTTTTATTTAGGTGTTTC-3’), pCMV-jT519D (5’-CCAAAACAAAAGAGATCAGGTGTTTCCACC-3’ and 5’-GGTGGAAACACCTGATCTCTTTTGTTTTGG-3’. All mutations were verified by DNA sequencing.

pRLSV40-Late was created as follows: the region of pSV010(-) (pUC18 containing the entire SV40 genome with a 4nt deletion at SV40 genome positions 5239-5242 in LT binding site II, rendering this plasmid inactive for SV40 DNA replication) corresponding to SV40 origin and promoter sequences nt 5097-272 was amplified using the following primers: 5’-TCACATGGCTCGACACCTTTCAAGACCTAGAAGGTCCATTAGCTG-3’ and 5’-GCACTGACTGCGTTACTGTGGAATGTGTGTCAGTTAGGGTGTGG-3’ (62,86). Site directed mutagenesis was then used to generate an NsiI restriction site at the 3’ end of the CMV promoter sequence of pRL using the following primer pair: 5’-GGTAGTTTATCACAGTTAAATGCATAACGCAGTCAGTGCTTCTG-3’ and 5’-CAGAAGCACTGACTGCGTTATGCATTTAACTGTGATAAACTACC-3’. This clone was digested with NsiI and BglII to remove the CMV promoter sequence and was combined with the pSV010(-) SV40 promoter fragment using the Infusion® HD Cloning kit (Clontech) such that the pSV010(-) late promoter sequence was cloned upstream of an intron (comprised of the 5’-region of a β-globin gene and 3’-region of an intron upstream of a heavy chain immunoglobulin gene included to promote splicing efficiency) and the Renilla luciferase gene both present in pRL (87). Clones were verified by DNA sequencing. Map of pRLSV40-Late generated by the Scalable Vector Graphics Plasmid Map tool found at Bioinformatics.org/savvy/.

γP^32^ATP-labelled 100mer helicase substrate was created as follows: 0.05 nanomoles of helicase substrate top strand (5’-AGTCAGAGACCCATAGAGACACACTTGCACAGTAACGCTTAGTCTCGTCAGATAGCAAGCCCAA TAGGAACCCATGTACCGTAACACTGAGTTTCGTCACTTTTTTTTTTTTTTTTTTTT-3’) were labelled with γP^32^ATP using T4 Polynucleotide Kinase according to the manufacturer’s protocol (Fermentas). End-labelled top strand was mixed with equimolar amount of bottom strand (5’-CCCCCCCCGTGACGAAACTCAGTGTTACGGTACATGGGTTCCTATTGGGCTTGCTAT CTGACGAGACTAAGCGTTACTGTGCAAGTGTGTCTCTATGGGTCTCTGACT-3’), placed at 95°C for 5 minutes, and the heating block was removed to room temperature to allow substrate to anneal slowly overnight. This creates a substrate with a 21nt unpaired polyT 3’ tail on the labeled strand and an 8nt unpaired polyC 5’ tail on the unlabeled strand. End-labelled, annealed substrate was purified with illustra ProbeQuant G-50 Micro Column (GE Healthcare). γ^32^P ATP-labelled 17mer helicase substrate was created by end labelling 0.05 nanomoles of a short oligo (5’-GTAAAACGACGGCCAGT-3’) with γP^32^ATP using T4 Polynucleotide Kinase according to the manufacturer’s protocol (Fermentas). End-labelled oligo was then mixed with equimolar amount of unlabeled, circular M13mp18 ssDNA (NEB), placed at 95°C for 5 minutes, and the heat block was removed to room temperature to allow substrate to anneal slowly overnight. End-labelled, annealed substrate was purified with illustra ProbeQuant G-50 Micro Column (GE Healthcare).

Poly dA/oligo dT substrate for Pol-Prim stimulation assays was created by mixing 100 nanomoles of poly [dA] (average length 250-500nt) (Roche) with 5 nanomoles of oligo [dT] (average length 18nt) (Roche) in TE (10mM Tris-HCl pH 8, 1mM EDTA) with 100mM NaCl in a 50μl volume. Mixture was placed at 95°C for 5 minutes and the heat block was removed to room temperature to allow substrate to anneal slowly overnight.

### Cell Culture

C33a, HEK293, and HEK293T cells were grown at 37°C with 5% CO_2_ in Dulbecco’s modified Eagle’s medium (DMEM) (Gibco) supplemented with 10% fetal bovine serum (Atlanta Biologicals) and 1% penicillin-streptomycin (Gibco). Sf9 insect cells were grown at 27°C in Grace’s Insect Cell Culture Medium (Gibco) supplemented with 4.175mM sodium bicarbonate, 10% fetal bovine serum, and 1% penicillin-streptomycin. High Five insect cells were grown at 27°C in Express Five Serum Fee Media supplemented with 16mM L-glutamine (Gibco) and 1% penicillin-streptomycin.

### Immuno-Blots

1 million C33a cells in a 10cm tissue culture dish were transfected with 1μg of pCMV-LT expression vectors for SV40 (WT, T518A, T518D) or JCPyV (WT, T519D) were lysed in 500μl of 1% Triton X-100 in PBS 48 hours post transfection. Lysate protein concentrations were normalized using a bicinchoninic acid BCA assay (Thermo Fisher) and run on 12.5% SDS-PAGE gels in sample buffer (50mM Tris-HCl pH 6.8, 100mM DTT, 2% SDS, 0.1% bromophenol blue, 10% glycerol) for 1.5h at 180 volts. Gels were transferred to a nitrocellulose membrane (GE) using a Novablot semi-dry transfer apparatus (Pharmacia Biotech) at 50 mAmp for 1h in transfer buffer (25mM Tris-HCl, 190mM glycine, 0.1% SDS, 20% methanol). Membranes were blocked for 1h at room temperature with 5% milk in TBST (50mM Tris-HCl, 150mM NaCl, 0.1% Triton-X100), washed 3 x 5min with TBST and placed in Ab101 (for SV40 LT, produced in house from pAB101 hybridoma cell culture and purified on protein A Sepharose (GE)) or Ab416 (for JCPyV LT, EMD Millipore) primary antibody diluted to 0.2μg/mL in TBST for 4h at RT. Membranes were washed 3 x 5min with TBST and placed in HrP-linked rabbit anti-mouse secondary antibody (Abcam) (Ab diluted to 0.5μg/mL in TBST for 2h at RT). Membranes were washed 3 x 5 min with TBST and developed using chemiluminescent substrate (Thermo Scientific SuperSignal West) and imaged using a BioRad Chemidoc.

### Computational prediction of phosphorylation sites on LT

NetPhos 3.1 Server was used to predict Ser, Thr, Tyr phosphorylation sites in the SV40 LT protein (54). The server uses the consensus phosphorylation sites of 17 protein kinases, including ATM (but not ATR), upon which the predictions are based. Similarities between the consensus sequences between ATM and ATR make ATM a reasonable exemplar for ATR.

### Immunoprecipitation of LT and Mass Spectrometry (MS)

HEK293T cells were grown to 70-90% confluence in complete DMEM with 10% FBS as described above. Cells were treated with 50lllM etoposide solution (Calbiochem) in DMSO (ETO) or only DMSO (Mock) for 2 hours at 37⁰C. Cell lysates were prepared using NP40 lysis buffer (ThermoFisher) containing 1% NP40, 250mM NaCl, 50mM Tris-Cl (pH 7.4) with protease and phosphatase inhibitors (Thermo Fisher #78440, 78420, Sigma # 45-P5726, 45-P0044), on ice, subjected to centrifugation in a microfuge for 15 min at 4C at 15.9K rcf. Protein levels were standardized using BCA assay (Thermo Fisher), flash frozen in liquid nitrogen and stored at -80⁰C. A 50 ul slurry of Dynabeads Protein G (Invitrogen, Carlsbad, CA) was used to immunoprecipitate LT using monoclonal Ab419 from ∼ 500ul ETO and DMSO extracts (1.3 mg/ml protein concentration). The proteins were eluted from the beads using NuPAGE LDS reducing sample buffer (ThermoFisher), boiled, resolved on 4-20% Novex Tris-glycine gels (ThermoFisher), and used for immuno-blotting (via iBlot 2 gel transfer using transfer template program P0) following standard procedures detailed above. Primary antibodies used were Ab101 (against large T-antigen) and Phospho-(Ser-Glu/Thr-Glu) ATM/ATR substrate antibody (pSQ/TQ) (Cell signaling Technology # 2851). Image J (NIH) was used to quantify protein bands.

LT-specific monoclonal antibody (100 ug of either Ab101 or Ab419 as designated, produced from hybridoma cell line supernatants and purified using protein A Sepharose) were bound to 5mg of M-280 Tosylactivated Dynabeads (Invitrogen # 14203) O/N at 37⁰C on a rotating wheel, washed in PBS and prepared according to manufacturer’s instructions. Subsequent binding reactions were assembled with 1mg beads in PBS mixed with 1.3mg mock or ETO-treated extracts at 3.0mg/ml final concentration for 4h at 4⁰C with tumbling. Beads were washed with PBS three times and eluted with 0.2% Formic acid 4X at RT. The elution was flash frozen in liquid N2 and dried in a speed-vac. Other immuno-precipitations were boiled in reducing SDS loading buffer at 95⁰C for 5 min, the bound proteins resolved using 4-20% Novex Tris-glycine gels, and silver-stained with MS-compatible Pierce Silver Stain for Mass Spectrometry (Thermo Scientific # 24600). The LT protein bands were excised from the gel, destained, and partially dried in 25mM ammonium bicarbonate in 50% acetonitrile. Trypsin digestions of both the solvent-elution and destained gel slices were used for mass spectrometric analysis at the Proteomics and Bioanalysis Core (PBC) Facility, New York State Center of Excellence in Bioinformatics and Life Sciences (NYS CoEBLS), Buffalo, NY.

Liquid chromatography-mass spectrometry (LC-MS) was carried out on a Dionex Ultimate 3000 nano LC system, a Dionex Ultimate 3000 gradient micro LC system with an WPS-3000 autosampler, and an Orbitrap Fusion Lumos mass spectrometer (ThermoFisher Scientific, San Jose, CA). (MS method details are included in the legend to Supplemental Figure 1.) Data Analysis: LC-MS rawfiles were searched by Sequest HT (embedded in Proteome Discoverer v1.4.1.14, Thermo Scientific) against SV40 LT sequence. The search parameters include: 1) Precursor ion mass tolerance: 20 ppm; 2) Fragment ion mass tolerance: 0.02 Da (OT)/0.8 Da (IT); 3) Maximal missed cleavages: 2; 4) Fixed modification: cysteine carbamidomethylation; 5) Dynamic modification: methionine oxidation, peptide N-terminal acetylation, serine/threonine/tyrosine phosphorylation; 6) Maximal modifications per peptide: 4. LT sequence coverage and peptide list was exported from Proteome Discoverer.

### Dual luciferase, cell-based transient viral DNA replication assays

SV40 and JCPyV cell-based DNA replication assays were performed as previously detailed (61,70). C33a cells were plated at a density of 25000 cells/well in white, flat bottom, 96 well plates (Corning) 20h prior to transfection. The transfection mixture for each well contained 100ul Opti-MEM (Gibco), 0.2ul lipofectamine (Invitrogen), 2.5ng of corresponding SV40 or JCPyV-origin containing Firefly Luciferase plasmid (pFLORI40 or pFLORIjc), 0.5ng of Renilla Luciferase plasmid lacking a viral origin (pRL) and increasing amounts of pCMV WT or mutant LT expression plasmid for SV40 or JCPyV. Transfection mixtures were replaced with fresh culture media 4h post-transfection. 72h post-transfection, Firefly and Renilla luciferase levels in cells were measured sequentially according to the Dual-Glo Luciferase assay protocol (Promega).

### Cell-based SV40 transcription activation assay

C33a cells were plated at a density of 25000 cells/well in white, flat bottom, 96 well plates 20h prior to transfection. Increasing amounts of pCMV-sWT or pCMV-sT518D were co-transfected into cells with 10ng plasmid pRLSV40-Late. 72 hours post transfection, Renilla luciferase levels were measured according to the Dual-Glo Luciferase assay protocol (Promega).

### Baculoviruses

p941-WT (a kind gift from Dr. Dan Simmons, University of Delaware) and p941T518D transfer vectors were combined with Sapphire Baculovirus DNA in sf9 insect cells according to the kit protocol (Allele Biotechnology) to generate WT LT and LT T518D-encoding baculoviruses, which were then further amplified according to the kit protocol.

### Infection, Expression and Purification of WT and LT T518D

Protocol adapted from Dr. Dan Simmons (88): T150 flasks with High Five insect cells at 90% confluency were infected with either WT LT or LT T518D baculoviruses at an MOI of three for 48h at 27°C. Infection media was removed by aspiration and 10mL of wash buffer TD (25mM Tris-HCl, 136mM NaCl, 5.7mM KCl, 0.7mM Na_2_HPO_4_, pH 7.4) was added to each flask, followed by cell scraping. Cells were pelleted in buffer TD at 570 RCF for 10 minutes at 4°C in a Beckman Coulter JS-4.2 rotor. Buffer TD was removed, and cells were lysed on ice for 30 minutes in 5mL Buffer B (150mM Tris-HCl, 150mM NaCl, 1mM EDTA, 10% glycerol, 0.5% NP40, 1mM PMSF, pH 8) per flask of cells. Lysates were centrifuged at 17500 RCF for 20 minutes at 4°C in a Beckman Coulter JA-20 rotor. Supernatants were incubated with 1mL of Protein A Sepharose (GE) saturated with and cross-linked to anti-LT antibody pAb101, for 2h on a shaker at 4°C. Beads were centrifuged for 2 minutes at 600 RCF at room temperature and the supernatant was removed. Beads were washed 5 times in 10mL buffer C (50mM Tris-HCl, 500mM LiCl_2_, 1mM EDTA, 10% glycerol, pH 8) followed by five washes in buffer D (10mM PIPES, 5mM NaCl, 1mM EDTA, 10% glycerol, pH 7.4). Beads were transferred to a small column at 4°C in buffer D. LT was eluted from beads 5 times with 0.5mL buffer E (20mM triethylamine, 10% glycerol, pH 10.8). Protein-containing fractions were pooled and dialyzed against buffer F (10mM PIPES, 5mM NaCl, 0.1mM EDTA, 1mM DTT, 10% glycerol, pH 7) for 16h at 4°C. Infection of twenty-five T150 flasks yields approximately 1mg of purified LT.

### *In vitro* SV40 DNA replication assays

SV40 *in vitro* DNA replication assays were performed as previously detailed (13). 60 ng of SV40-origin template pSV011 (or pSV011(-), a template with a 4nt deletion that renders the SV40 origin non-functional) was incubated with increasing amounts of purified WT or T518D SV40 LT, in replication buffer (30mM Tris-HCl pH7.5, 40mM creatine phosphate, 7mM MgCl_2_, 4mM ATP, 200μM CTP, 200μM GTP, 200μM UTP, 100μM dCTP, 100μM dGTP, 100μM dTTP, 25μM dATP, 0.5M DTT), 0.1mg/ml acetylated BSA, 1μCi αP^32^dATP, and 40ug of HEK293 cyto/nucleosolic extract in 10μl volume. Reactions were incubated at 37°C for 1h and terminated by addition of 10μl stop buffer (20mM Tris-HCl pH7.5, 10mM EDTA, 0.1% SDS, 1μg/μl proteinase K) for 20 minutes at 37°C. Reactions were quantified by spotting on DE81 filter paper (Whatman), washing away the unincorporated nucleotides with 0.5M NH4PO4, and scintillation counting. DNA products were extracted using phenol:chloroform, precipitated using ammonium acetate and ethanol, and resolved on 0.8% agarose, Tris/borate/EDTA gels. Gels were fixed for 10 minutes in 15% methanol/15% acetic acid, rinsed in ddH_2_O, dried, and analyzed by phosphorimager autoradiography using a Molecular Dynamics Typhoon system.

### Purification of human Topoisomerase I, Replication Protein A, and Polymerase-α/Primase

Full length human Topoisomerase I (Topo I) was purified from extracts of baculovirus-infected Hi-5 cells using Mono Q, Mono S, and POROS columns as described previously (89). Full-length recombinant human replication protein A (RPA) was expressed in *Escherichia coli* and purified as described previously (90). Human Polymerase-α/Primase (Pol-Prim) was purified from human 293 cell extracts using immuno-affinity chromatography using an anti-Pol-Prim antibody mAB1645 Sepharose column as described previously (91).

### Enzyme Linked Immunosorbent Assays (ELISAs)

*For RPA and TopoI*: Dilutions of RPA or TopoI were prepared in 50μl TBS (25mM Tris-HCl pH 7.5, 150mM NaCl) were immobilized in each experimental well of vinyl, 96-well ELISA plate (GIBCO) by rocking for 1h at RT. Wells were washed 3X with TBST (TBS with 0.1% Triton X100), blocked with 200μl TBST +1% dry milk/1% BSA for 1h at RT with rocking, and washed 3 more times with TBST. Dilutions of increasing amounts of WT or LT T518D were prepared in 50μl TBS, added to the appropriate wells, and incubated for 1h at RT with rocking. For block controls, the highest amount of WT LT was added to blocked wells to determine non-specific interaction background. All wells were washed 3X with TBST. 50μl of SV40 LT pAb101 primary antibody, diluted to 0.2μg/ml in TBST, was added to each well and incubated at RT for 1h with rocking. Wells were washed 3X with TBST. 50ul of horseradish peroxidase (HrP)-linked anti-mouse secondary antibody, diluted 1:5000 in TBST, was added to each well followed by incubation for 1h at RT with rocking. Wells were washed 7X with TBST, followed by 3X with TBS. Wells were incubated with 50μl HrP substrate solution (for 1mL of fresh solution, combine 100μl 1.1M sodium acetate pH5.5, 16.7μl of 25mM 33’55’tetramethylbenzidine, 0.3μl 30% H_2_O_2_, and 883μl ddH_2_0) for 10 min at RT. 50ul of stop solution (2M sulfuric acid) was added to each well and plate was read on microplate reader at absorbance 450. *For Polymerase α/Primase*: protein incubation order was reversed; 50ng of either WT or T518D was immobilized as first protein, dilutions containing increasing amounts of Pol-Prim were added as second protein, primary antibody was 0.2μg/ml anti-Pol-Prim pAB1645 (produced from pAB1645 hybridoma cell culture), and secondary antibody was 0.2μg/ml HrP-linked rabbit anti-mouse antibody (Abcam). Wash and development steps were the same as for the RPA/TopoI ELISAs. The highest amount of Pol-Prim was added to blocked wells to determine non-specific background.

### Poly dA/oligo dT Polymerase-α/Primase Stimulation Assays

Increasing amounts of purified WT or LT T518D was incubated with 50ng purified Pol-Prim, 1μCi αP^32^dTTP, reaction buffer (40mM Tris-HCl pH 6.9, 6mM MgCl_2_, 40μM dTTP, 10% glycerol, 1mM DTT, 40μg/mL BSA), and 0.04 nanomoles of poly[dA]/oligo[dT] substrate in 10μl reactions for 45 minutes at 37°C. Reactions were terminated by addition of 10μl stop buffer (0.5M NaOH, 7mM EDTA, 4% Ficoll) and 2μl 0.2% Bromophenol Blue. Reactions were quantified by spotting on DE81 filter paper (Whatman), washing away unincorporated nucleotides with NH4PO4, followed by scintillation counting. DNA replication products were resolved on agarose gels poured at 1% agarose/1mM EDTA/30mM NaCl (first equilibrated in alkaline-denaturing buffer for 2 hours) and run in alkaline-denaturing buffer (50mM NaOH, 1mM EDTA) at 6 volts/cm for 4h at RT. Gels were fixed for 10 minutes in 2 volumes 15% methanol/15% acetic acid, rinsed in ddH_2_0, dried, and analyzed using a Molecular Dynamics Typhoon system.

### ATPase Assays

250ng purified WT or LT T518D was combined with 1μCi γP^32^ATP in reaction buffer (30mM Tris-HCl pH 7.8, 7mM MgCl_2_, 0.1mM ATP pH 7.5, and 0.5mM DTT) in 10μl. Time course reactions were carried out at 37°C as indicated. Increasing amounts of poly(dT) (average length 221bp) (Bioline) or additional unlabeled ATP (NEB) were added to each reaction as indicated. Poly(dT) and cold ATP titration reactions were incubated at 37°C for 30 minutes. All ATPase reactions were stopped by addition of 0.5μl of 0.5M EDTA. 1ul of each reaction was spotted 2cm from the bottom of a thin layer polyethyleneimine chromatography (TLC) plate (Sigma Aldrich). TLC plate was developed in 0.5M LiCl_2_/1M formic acid until buffer front was 2cm from top of plate. TLC plates were dried and analyzed using a Molecular Dynamics Typhoon system. Density of spots containing γPO_4_^3-^ hydrolyzed during the ATPase reactions were quantified using ImageJ software.

### DNA Helicase Assays

250ng purified WT or LT T518D was incubated with 50 femtomoles of γP^32^ATP-labelled 17mer or 100mer helicase substrate and reaction buffer (30mM HEPES pH 7.5, 7mM MgCl_2_, 0.5mM DTT, 4mM ATP, 0.1mg/ml BSA) in 10μl volumes at different points over a time course. Reactions were terminated by addition of 3ul of stop buffer (5% SDS, 50mM EDTA, 0.5mg/ml proteinase K, 12.5% glycerol, 0.25% bromophenol blue) at 37°C for 10 minutes. 10μl of each reaction was loaded on a 12% native TBE PAGE gel and run at 100 volts for 1.5h (until dye front reaches bottom of the gel). Gel was fixed in 15% methanol/15% acetic acid, rinsed in ddH_2_O, dried, and analyzed by phosphorimager autoradiography using a Molecular Dynamics Typhoon system. Images were quantified with ImageJ software.

## Acknowledgements

We thank the following for their generous gifts: especially Dan Simmons (for p941-WT and numerous other reagents, advice and protocols), Amelie Fradet-Turcotte and Jacques Archambault (pFLORI40, pCMV-sWT, pRL), and Peter Bullock and J. Archambault (pFLORIjc and pCMV-jWT). We also thank J. Archambault and Mark Sutton for valuable advice and suggestions throughout this study, Jeffrey C. Martin for assistance in editing of the manuscript. Dr. Wei Yang (NIH/NIDDK) for helpful discussion and advice regarding SV40 LT structure and function and the predicted effects of LT phosphorylation and the mutations evaluated in this manuscript.

## Figures

**Supplemental Figure 1.**
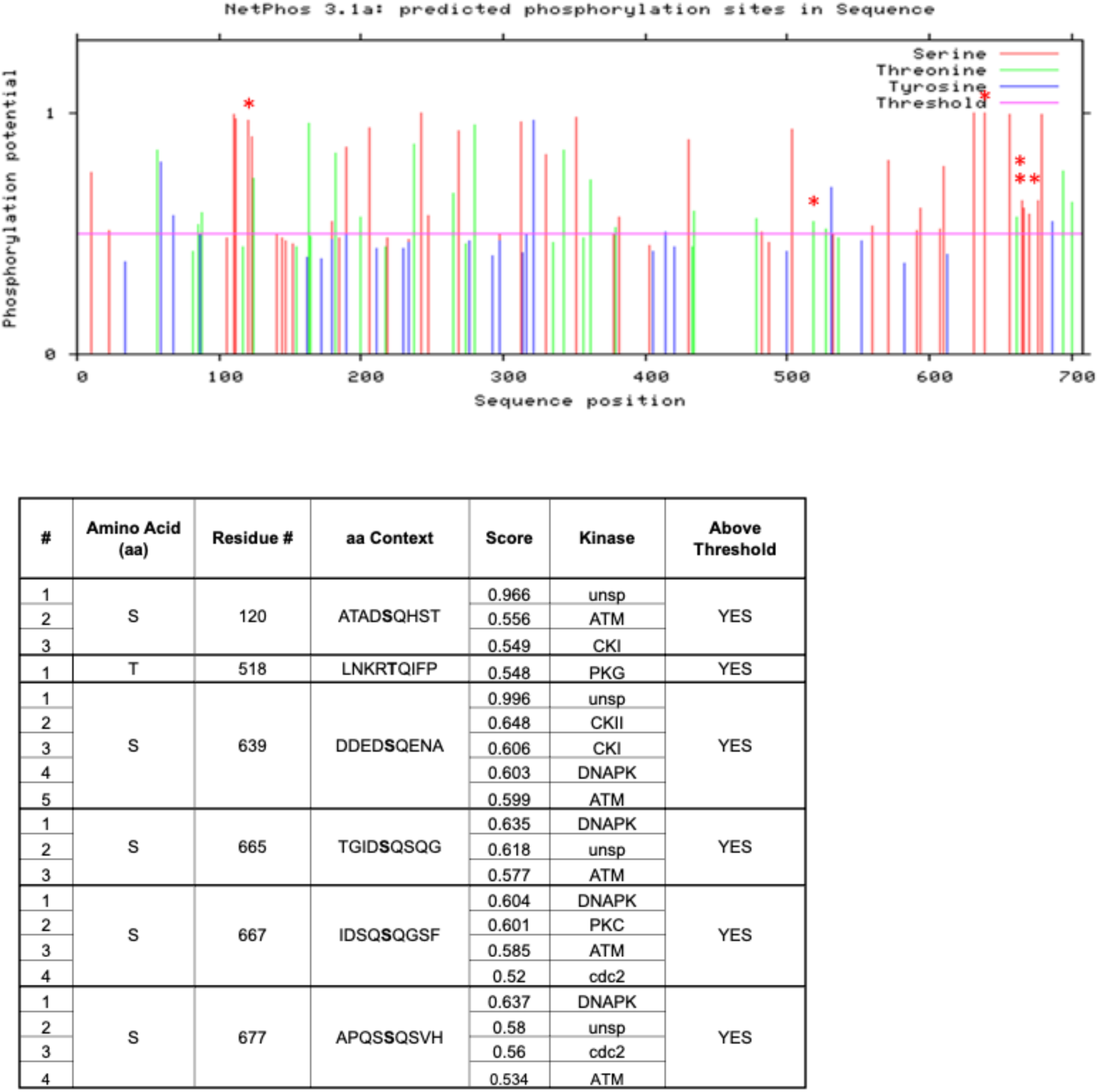
NetPhos 3.1-predicted phosphorylation sites on SV40 LT. SV40 LT sequence was submitted to computational analysis via the NetPhos 3.1 server. The top panel is the NetPhos3.1a readout for LT, showing all S, T, and Y residues. The axis represents the calculated potential phosphorylation of the indicated aa residue, and the ordinate represents the linear aa sequence of LT. The pink Threshold line at 0.500 represents a value for which if aa residues have a calculated Phosphorylation potential above this line, these residues are predicted to be phosphorylated in vivo. The red asterisks are directly above the residues indicated in the lower panel. The lower panel is the NetPhos3.1 results readout for SV40-LT, showing the S and T residues (bold) of the conserved six SQ/TQ motifs in context of +/- 4 amino acids. The linear sequence motifs surrounding phosphorylated residues have been utilized with 17 known kinase–substrate specificities. The score for each of the S and T residues predicted to be a phosphorylation site (YES) is calculated. All positive predictions with a score higher than the threshold value are included and all negative predictions with scores lesser than the threshold value are excluded.

**Supplemental Figure 2.**
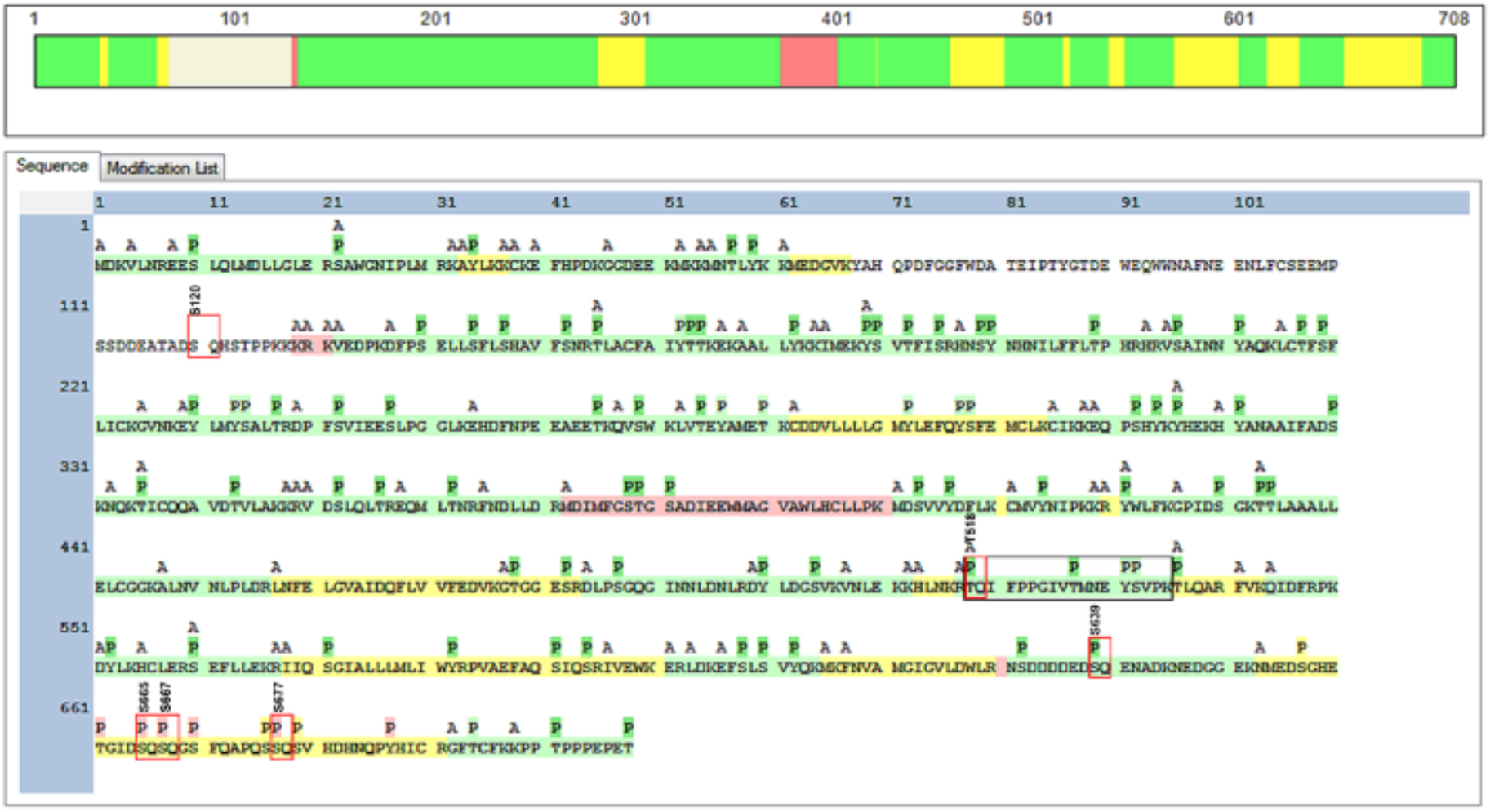
Map of phospho-sites on SV40 LT using MS2 analysis. Sequence coverage of SV40 large T antigen (LT). Identified peptides and S/T/Y phospho-sites (P) are highlighted with different colors denoting high (green), medium (yellow) and low (red) confidence levels. Over 90% of the protein sequence was covered by MS analysis with 91 potential phospho-sites. Five of the six potential SQ/TQ phospho-sites (S120, T518, S639, S665, S667 and S677: red boxes) were localized with varying levels of confidence. A high confidence localization of T(518)Q phosphorylation was indicated by the analysis. The N-terminal acetylations (A) are marked. The previously demonstrated phosphorylation at S120 occurred within the small percentage of LT that was not covered by the MS analysis. For LT gel samples, an in-gel digestion protocol was employed. gel slices were first chopped into 1 cm^3^ cubes using a clean scalpel. Gel cubes were transferred to new tubes, and were washed by 500μL ddH_2_O and then 1 mL acetonitrile (ACN; Each wash step was accompanied by 10 min constant shaking at room temperature). A volume of 1 mL 50% Tris-formic acid (FA)/ACN was then added and the gel cubes were incubated under 4°C overnight for destaining. Destained gel cubes were washed by 1mL water and three times of 1mL ACN, and protein was reduced by 500μL 10mM dithiothreitol (DTT) for 45 min in a covered thermomixer (Eppendorf, Hauppauge, NY) under 37°C with constant shaking. After washing with 1mL ACN, protein was alkylated by 500μL 25mM iodoacetamide (IAM) was added for protein alkylation for 45 min in a covered thermomixer under 37°C with constant shaking. Gel cubes were then washed by 1mL ACN for three times and completely dried. A volume of 300μL trypsin (Sigma-Aldrich, St. Louis, MO) dissolved in 50mM Tris-FA (0.0125μg/μL) was added to each sample, and the samples were put on ice for 30 min. Excess trypsin was removed and 200μL 50mM Tris-FA was added for overnight trypsinization under 37°C in a thermomixer with constant shaking. Trypsinization was terminated by addition of 20μL 5% FA with 15-min shaking, and supernatant was collected to new Eppendorf tubes. A volume of 100μL 50% Tris-FA/ACN was added to the remaining gel cubes with 15-min shaking, and the supernatant was combined with the previous one. Derived peptide mixture was dried using a SpeedVac and reconstituted by 50μL 0.1% trifluoroacetic acid (TFA)/1% ACN in ddH_2_O. Samples were centrifuged at 18,000 g under 4°C for 30 min and transferred to vials for analysis. For lyophilized LT samples, 50μL 0.5% SDS was added to each sample, and samples were sonicated for 30 sec and vortex for 10 min to reconstitute protein. Protein reduction and alkylation was performed sequentially by addition of 2μL 200μM DTT and 4μL IAM, each with 45-min incubation in a covered thermomixer under 37°C with constant shaking. Protein was then precipitated by two-step addition of 60 and 300μL chilled acetone, incubated under -20°C for 3hr, and centrifuged for 30 min at 18,000 g under 4°C to pellet protein precipitated. Pelleted protein was gently rinsed with 400μL methanol, decanted, and wetted by 45μL 50mM Tris-FA. A volume of 5μL trypsin dissolved in 50mM Tris-FA (0.25μg/μL) was added to each sample, and trypsinization was performed under 37°C overnight (∼16hr) with constant shaking in a covered thermomixer. Trypsinization was terminated by addition of 0.5μL FA, and derived peptide mixture was centrifuged at 18,000 g under 4°C for 30 min, and supernatant was transferred to vials for analysis. A single injection of 4 μL derived peptides were analyzed for each sample. The liquid chromatography-mass spectrometry (LC-MS) system consists of a Dionex Ultimate 3000 nano LC system, a Dionex Ultimate 3000 gradient micro LC system with an WPS-3000 autosampler, and an Orbitrap Fusion Lumos mass spectrometer (ThermoFisher Scientific, San Jose, CA). A large-i.d. trapping column (300µm ID × 5 mm) was implemented prior to nano LC column (75-μm ID × 100 cm, packed with 3μm Pepmap C18) separation for high-capacity sample loading, matrix component removal, and selective peptide delivery. Mobile phase A and B were 0.1% FA in 2% acetonitrile and 0.1% FA in 88% acetonitrile. The 180-min LC gradient profile was: 4−13% B for 15 min; 13% to 28% B for 110 min; 28% to 44% B for 5 min; 44% to 60% B for 5 min; 60% to 97% B for 1 min, and isocratic at 97% B for 17 min. MS was operated under data-dependent acquisition (DDA) mode, with a maximal duty cycle time of 3 sec. MS1 spectra were acquired in the m/z range 400∼1,500 under 120k resolution with dynamic exclusion settings (60 sec ± 10 ppm). Precursor ions were filtered by quadrupole using a 1-Th wide window and fragmented by high energy C-trap dissociation (HCD) at a normalized collision energy of 35%. MS2 spectra were acquired under 15k resolution in either Orbitrap or Ion Trap.

**Supplemental Figure 3.**
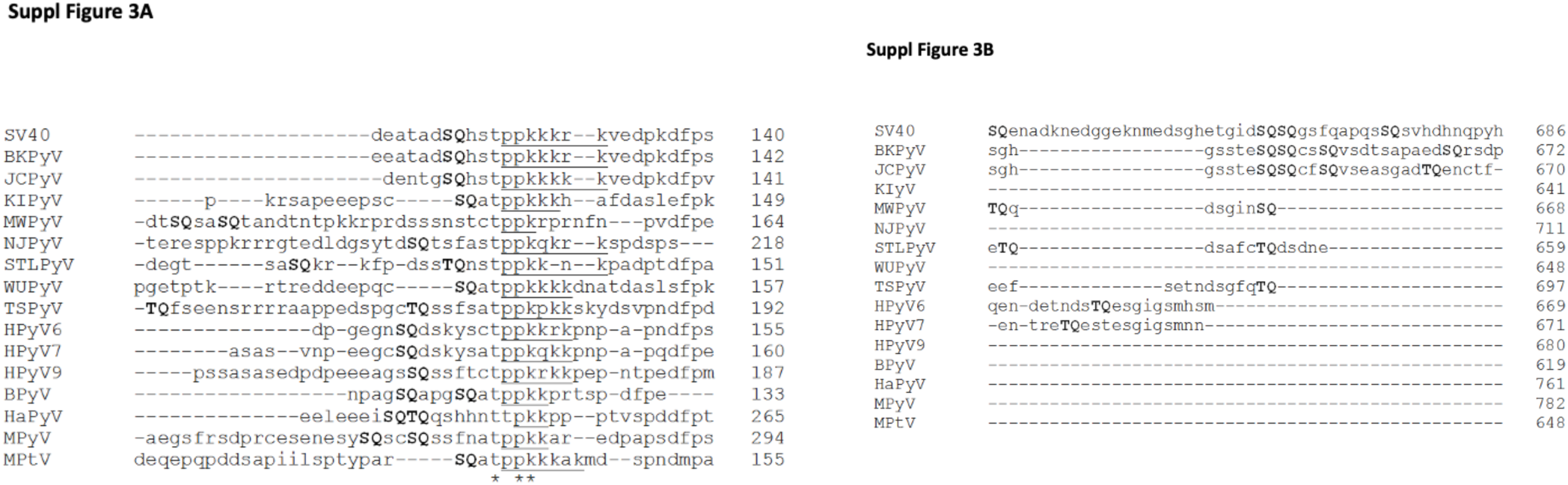
Amino Acid Alignment of Sixteen Mammalian Polyomavirus LTs Identifies Three Regions of SQ/TQ Sites. LT sequences from 12 human polyomaviruses (SV40, BKPyV, JCPyV, KIPyV, MWPyV, NJPyV, STLPyV, WUPyV, TSPyV, HPyV6, HPyV7, and HPyV10), bovine polyomavirus (BPyV), hamster polyomavirus (HaPyV), murine polyomavirus (MPyV), and murine pneumotropic virus (MPtV) identify three regions containing potential DDR Kinase substrate (SQ/TQ) sites (bolded): (A) a region just upstream of the known or predicted LT nuclear localization sequences (underlined), (B) a cluster of sites corresponding to the extreme C terminus of SV40 LT, and (Fig 1F) a region within the AAA+ ATPase/Helicase domain.

**Supplemental Figure 4.**
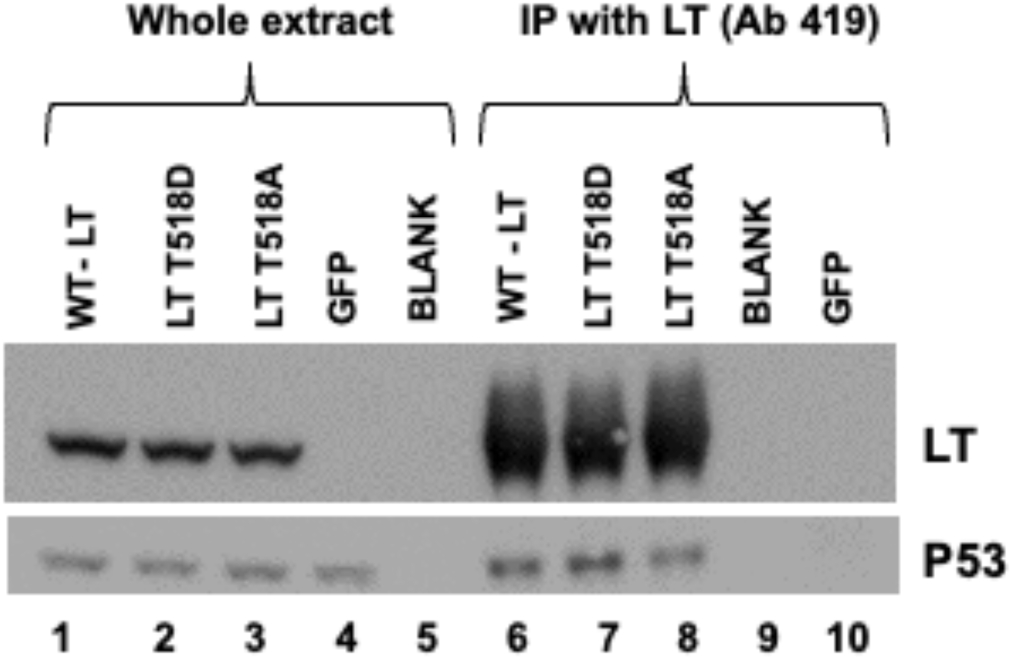
Phosphomimetic and non-phosphorylatable LT T519 mutations are as efficient as wt LT for co-immunoprecipitation of cellular p53. HEK293 cells were transfected with LT-WT, T518D, T518A. Whole protein extracts were subjected to immunoprecipitation of SV40 LT using monoclonal antibody Ab 419. SDS-PAGE was performed and immunoblotted with antibodies against LT (Ab 101) and p53 as described in Methods. LT and p53 was observed in the whole extracts (Lanes 1-3) of LT-WT, T518D, T518A transfected cells. Lanes 6-8 (IP with Ab419) showed an equivalent enrichment of LT from all three LT-WT, T518D, T518A transfected cell extracts along with co-precipitated p53. While p53 is clearly observed in the negative control (GFP transfected) whole cell extract (Lane 4), both LT and p53 are absent in the IP lane 10. This demonstrates that T518D or T518A mutations within the helicase region of SV40-LT does not affect binding to p53.

**Supplemental Figure 5.**
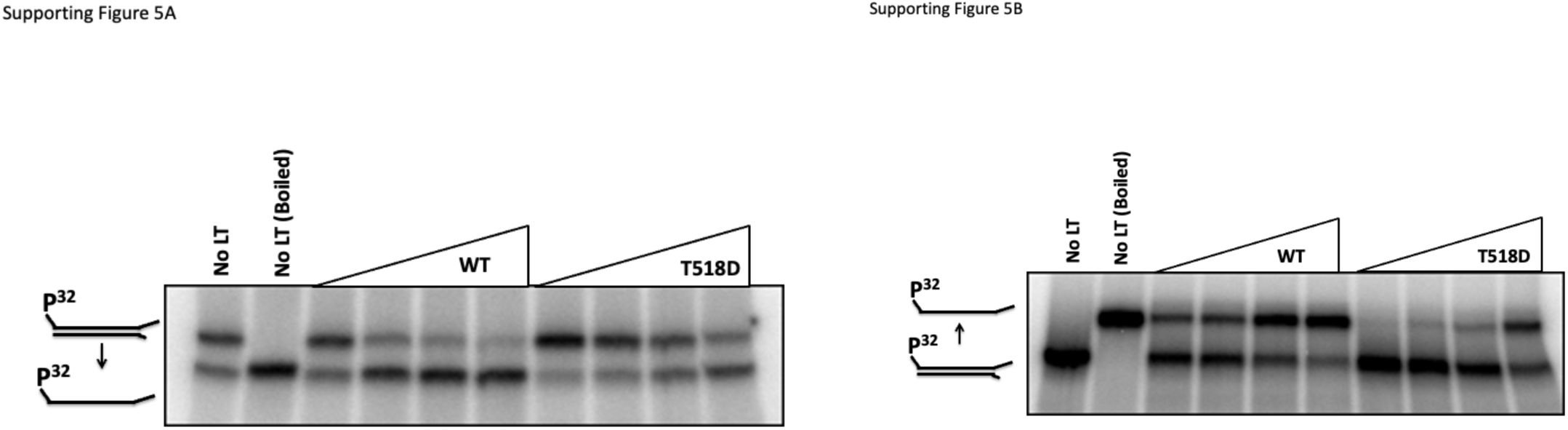
Mutant LT T518D is deficient for DNA Helicase Activity on both 50 and 100bp dsDNA substrates *in vitro*. Increasing amounts (125, 250, 500, or 1000ng as indicated by increasing triangles) of WT or LT T518D were incubated with (A) a 50bp or (B) a 100bp dsDNA helicase substrate with a single P^32^ end-labelled strand. Reactions were resolved on native TBE gels, exposed to phorphor imaging-screens, and analyzed by autoradiography. Values were obtained by setting the density of the released, radiolabelled band of a boiled reaction to 100% helicase activity and comparing densities of released bands in reactions to the boiled controls. (A) and (B) are representative images for 50mer and 100mer LT-titration helicase assays. “No LT” controls (lanes 1) show migration of intact, duplex DNA helicase substrates compared to “No LT (Boiled)” controls (lanes 2) showing migration of the single, radio-labelled strand of the helicase substrate following separation of DNA strands (note substrate cartoons highlighting the difference in direction of migration of the boiled 50mer and 100mer substrates). Note the small amount of end-labelled, unannealed ssDNA present in the “No LT” lane 1 control of panel A.

